# Development positions malignant cellular states but does not explain their diversification

**DOI:** 10.64898/2026.05.28.728220

**Authors:** Juan F Poyatos

## Abstract

Epithelial cancers are often described as aberrant reactivations of embryonic or tissue-forming programs, but whether malignant cellular-state diversification is actually constrained by developmental trajectories remains unclear. Here, we present a quantitative framework to test this idea in pancreatic ductal adenocarcinoma (PDAC). Using representation learning on large-scale single-cell data, we build a reference space that captures the main axes of normal foregut and pancreatic epithelial variation. Malignant cells can be mapped into this space, showing that developmental biology helps interpret their identity. However, these coordinates account for only part of the variation. A substantial portion lay outside the reference space, and the residual component was structured rather than diffuse. Furthermore, displacements between cancer states showed weak alignment with canonical axes and did not correspond to simple progression along them, even when they retained measurable components within the broader subspace. Thus, developmental programs offer a useful coordinate system for describing where PDAC states sit relative to normal epithelia, but they do not explain the directions along which malignant states differ. Instead, malignant diversification follows additional structured, cancer-adaptive axes outside the dominant geometry of normal development. More generally, these results demonstrate how disease-state variation can be decomposed into shared coordinates of normal identity and distinct directions of pathological reprogramming.

**Significance Statement:** Tumors are often said to reuse genetic programs from early tissue formation, but similarity to those programs does not mean cancer cells follow the same trajectories. We developed a quantitative framework to separate these ideas in pancreatic ductal adenocarcinoma. Variation tied to normal epithelial differentiation provided a useful coordinate system for locating malignant cell states, yet differences between those states showed weak alignment with canonical differentiation directions and included structured, cancer-adaptive programs. Thus, pancreatic cancer retains aspects of normal cell identity while diversifying along additional malignant axes. This distinction clarifies how insights from normal tissue programs should be applied to cancer plasticity and provides a general method for comparing disease states with normal cellular programs.

## Introduction

Cancer progression is shaped not only by genetic diversification but also by the emergence of heterogeneous cellular states (Fidler 1978; Torborg et al. 2022; Feinberg et al. 2023). Early studies of tumor evolution focused on mutational lineages and clonal expansion (Greaves et al. 2012; Shah et al. 2012; Frankell et al. 2023), but single-cell profiling has revealed that tumors occupy complex phenotypic landscapes (Marjanovic et al. 2020; Burdziak et al. 2023). Within a single malignancy, cells can adopt contrasting transcriptional programs, including stem-like, progenitor-like, inflammatory, stress-responsive, epithelial–mesenchymal, and lineage-divergent states (Tirosh et al. 2024). This phenotypic variety is increasingly recognized as a driver of adaptation, metastasis, and therapeutic resistance (Marine et al. 2020; Househam et al. 2022).

A central question is how this diversity is organized. Do cancer cells explore state space freely, or are their identities constrained by pre-existing biological organization? Hierarchical models emphasize stem-like populations that generate downstream phenotypic mixture (Batlle et al. 2017), whereas other models suggest that cancer cells continuously sample a high-dimensional state space with relatively weak intrinsic constraints (Shaffer et al. 2017). A third view proposes that malignant cell states are shaped by developmental programs of the tissue of origin (Neftel et al. 2019; Tirosh et al. 2024; Patel et al. 2024). This constraint model provides a powerful explanation for the recurrence of related cell states across tumors. However, it also raises a more specific question: does developmental structure simply position malignant states, or does it constrain the directions along which they differ?

This distinction matters because resemblance is not equivalent to trajectory. Development may provide a coordinate system for interpreting cancer cell identity without defining the dominant directions of variation. Existing marker-based analyses, lineage annotations, and trajectory-inference methods can identify similarities between cancer and development, but they often use the same developmental framework both to position malignant states and to interpret differences among them (Trapnell et al. 2014; Street et al. 2018; Stuart et al. 2019). What has been missing is a quantitative approach that separates these two questions.

Here, we address this problem in pancreatic ductal adenocarcinoma (PDAC), a malignancy characterized by extensive epithelial plasticity (Burdziak et al. 2023). PDAC is a particularly relevant setting because recent work has emphasized plasticity as a driver of its dissemination and treatment resistance (Jiménez-Sánchez et al. 2026; Innamorati et al. 2026; Gautam et al. 2026). These observations raise a more precise question: are plastic tumor states organized by developmental trajectories, or do they only retain some developmental features while diversifying along cancer-adaptive axes?

We use a masked language model (MLM), a representation-learning approach adapted from natural language processing, to learn context-aware cellular embeddings from large single-cell datasets (Devlin et al. 2019; Yang et al. 2022; Theodoris et al. 2023; Cui et al. 2024; Szalata et al. 2024; Poyatos 2025). This choice is central to the analysis: the question is not only whether cancer cells express selected developmental markers, but whether they share *distributed* transcriptional structure with normal states. By training the model to predict masked genes from the remaining transcriptional context, the MLM learns relationships among genes and cell states that may not be captured by marker-based analyses or purely linear decompositions of gene-expression space. Developmental annotations and malignant-state labels are then used after embedding to define reference spaces, lineage directions, and state summaries.

We construct a data-driven reference space of normal foregut and pancreatic epithelial development (Qiu et al. 2024), defining both a developmental PC subspace and lineage directions from foregut epithelium toward ductal, acinar, and biliary states. We then embed PDAC cells (Burdziak et al. 2023) in this framework and decompose malignant variation into developmental and residual components. This separates positional alignment, which asks where cancer states lie relative to developmental identities, from directional alignment, which asks whether differences between states follow developmental directions. Our analysis shows that these two aspects are decoupled. Malignant states can be positioned relative to developmental geometry, but their differences show weak alignment with canonical lineage axes and retain substantial residual components. Thus, developmental genetic programs provide a useful coordinate system for interpreting PDAC cell identity, but they do not define the dominant directions of variation. More broadly, this work provides a framework for comparing disease states with normal biology by separating shared coordinates of identity from disease-associated axes of variation.

## Results

### Development defines a low-dimensional reference space that does not capture most malignant variation

To define a quantitative reference for normal epithelial variation, we analyzed a mouse prenatal single-cell dataset spanning approximately from embryonic day 8.5 to postnatal day 0 (E8.5 to P0) (Qiu et al. 2024). Cells were represented by classification-token (CLS) embeddings from a masked language model (MLM) (Devlin et al. 2019) trained jointly on this atlas and a mouse PDAC model cohort (Burdziak et al. 2023) (Fig. 1A; Methods; Supplement; Figs. S1–S7). We first performed principal component (PC) analysis on developmental cells (Jolliffe 2002). The resulting PCs define a *reference* coordinate system for the dominant modes of normal epithelial variation.

**Figure 1:**
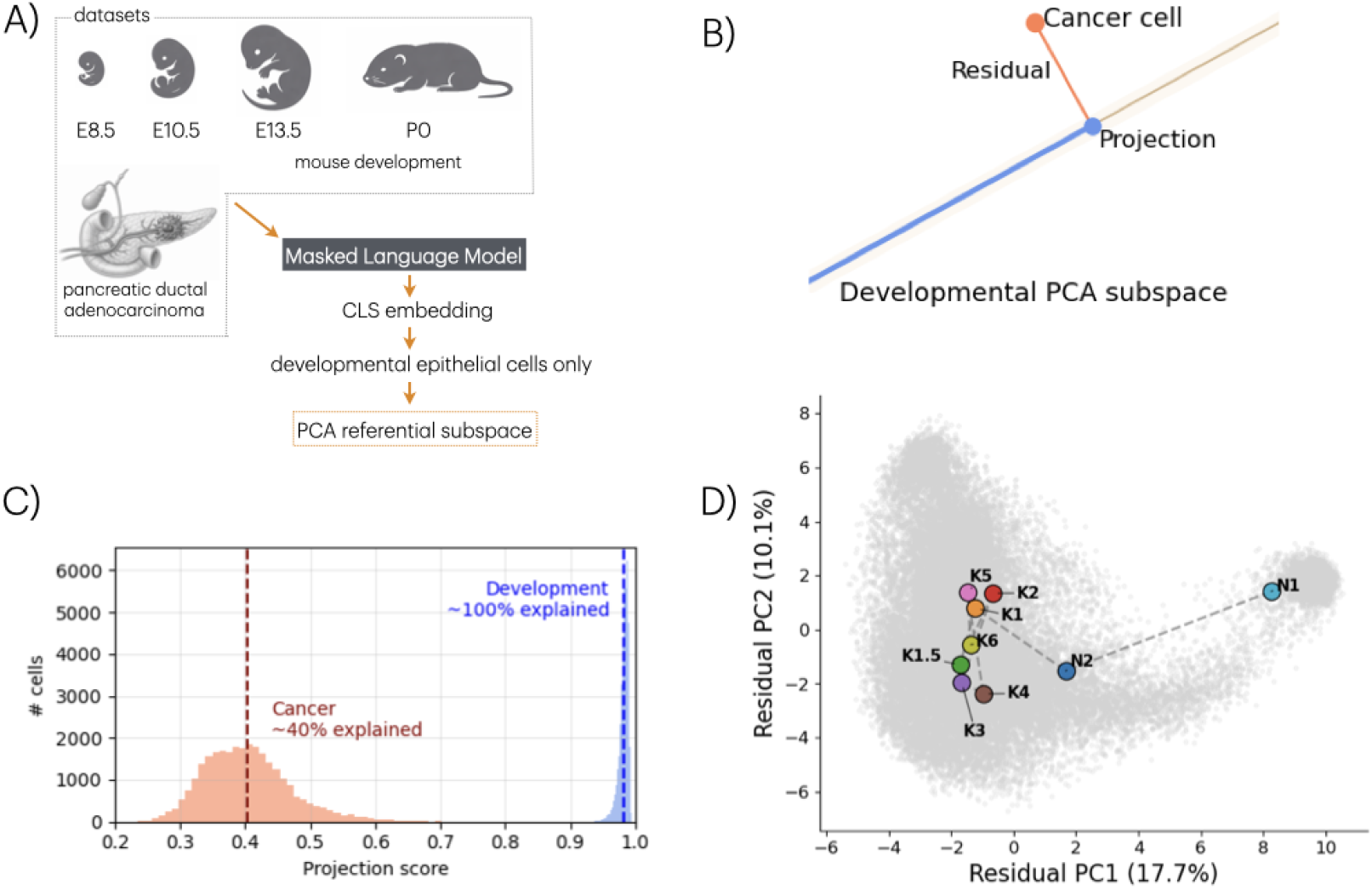
Developmental variation defines a low-dimensional reference space that captures only part of malignant-cell variation. A) Mouse prenatal developmental epithelial cells, from approximately embryonic day 8.5 (E8.5) to postnatal day 0 (P0), and pancreatic ductal adenocarcinoma (PDAC) epithelial cells were jointly embedded with a masked language model, and classification-token (CLS) embeddings from developmental epithelial cells were used to define a reference space for normal epithelial variation with principal-component analysis (PCA). B) Each cancer-cell embedding was decomposed into two parts: a component reconstructed within the developmental PCA subspace and a residual component orthogonal to that subspace. C) *Per*-cell projection scores, defined as the fraction of embedding magnitude captured by the developmental reference space. Developmental cells have scores near one, as expected because this space was learned from developmental variation. In contrast, PDAC cells have substantially lower scores, indicating that much of their variation lies outside dominant developmental axes. D) Cancer cells projected onto the first two PCs of the residual embeddings. Condition centroids, including healthy pancreas N1, reversible metaplasia N2, preneoplasia K1–K2, pancreatic intraepithelial neoplasia K3–K4, malignant PDAC K5, and distal metastases K6, occupy distinct regions. Thus, the residual component is structured rather than isotropic noise.

The embeddings were highly structured. The top 75 PCs explained approximately 98% of variance, indicating that normal states occupy a relatively low-dimensional region within the 256-dimensional latent space (Supplement). To test whether this structure was biologically interpretable rather than only statistically compact, we annotated the leading PCs by correlating cell coordinates with individual gene expression and gene-program scores. These PCs recovered expected epithelial programs, including proliferative progenitor-like states and acinar/secretory programs (Supplement, Fig. S8). Thus, the developmental PC subspace provides a biologically interpretable coordinate system for normal variation.

How much heterogeneity among malignant cells can this reference space represent? For each cell, we decomposed its embedding into a component reconstructed from the developmental PC basis and a residual component orthogonal to that basis (Fig. 1B; Methods; Supplement). We summarized this decomposition using a *per*-cell projection score, defined as the fraction of the cell’s embedding magnitude captured by the PC subspace. Developmental cells had projection scores close to one, as expected because the space was defined from those cells (Fig. 1C). In contrast, PDAC cells had substantially lower scores, with a mean of approximately 0.4. Thus, only a minority of malignant embedding magnitude is explained by the dominant axes of variation.

The residual component was structured rather than random. PC analysis of the orthogonal residual embeddings showed that the top 30 residual PCs captured approximately 88% of variance (Fig. 1D; Supplement), indicating that tumor cells contain additional low-dimensional organization outside the reference space. Consistent with this interpretation, both the developmental projection and residual component carried information for discriminating cancer cell states, with modest additional predictive performance when the two components were combined (Supplement).

We then examined the biological programs encoded by the malignant residual component (Supplement, Fig. S9). The leading residual PCs captured coherent cancer-associated programs (Barkley et al. 2022; Tirosh et al. 2024), including acinar/secretory differentiation, epithelial-to-mesenchymal transition (EMT) and extracellular-matrix (ECM) remodeling, AP-1 immediate-early response signaling, glycolytic and mitochondrial metabolic rewiring, hypoxia, oxidative stress, KRAS/MAPK signaling, stemness or progenitor-like identity, MYC-associated proliferation, and unfolded protein response programs. For example, the acinar/secretory loading of residual PC1 was marked by digestive enzyme and secretory genes, including Pnlip, Ctrc, Pla2g1b, Zg16, Cela1, Ctrl, and Aqp12. Residual PC2 captured a prominent EMT/ECM-remodeling and stress-adaptive invasive program. These results show that the residual is not merely unexplained technical variance, but contains structured cancer-associated biology not captured by the primary developmental basis.

### Malignant-state differences are weakly aligned with canonical developmental directions

We next focus on whether differences between malignant states align with the directional organization of *normal* epithelial differentiation. We first defined a lineage-referenced co-ordinate system using centroid-to-centroid vectors from foregut epithelium toward three terminal or adjacent epithelial states: ductal, acinar, and biliary cells (Gittes 2009; Shih et al. 2013) (Methods; Supplement). These directions were separated by pairwise angles of approximately 54–60^*°*^, indicating that epithelial differentiation is organized across multiple directions rather than along a single dominant axis (Supplement, Fig. S6). Developmental cells populated broad corridors around these directions, consistent with a continuous and overlapping manifold rather than sharply separated one-dimensional trajectories (Supple-ment, Fig. S7).

We then computed centroid-to-centroid displacement vectors between annotated cancer states and compared them with the previous directions. Following the original annotations (Burdziak et al. 2023), we considered three malignant states: ductal, transitional, and malignant ductal. These states define three displacements: `early`, from ductal to transitional; `late`, from transitional to malignant ductal; and `total`, from ductal to malignant ductal. These vectors quantify differences between annotated states and should not be interpreted as directly observed temporal trajectories.

Malignant displacement vectors showed weak alignment with all three canonical axes. Across comparisons, cosine similarities were small in magnitude (Fig. 2A), indicating that the deviations are not organized as simple movement along developmental directions. We then asked whether these same state differences were nevertheless partly captured by the full subspace. To test this, we decomposed each displacement vector into a component reconstructed within the 75-dimensional PC subspace and an orthogonal residual component. A measurable fraction of each displacement lay within the subspace, but substantial residual components remained outside it (Fig. 2B; Supplement). Thus, cancer states are not developmentally disconnected, but their differences are not reducible to canonical developmental branch geometry.

**Figure 2:**
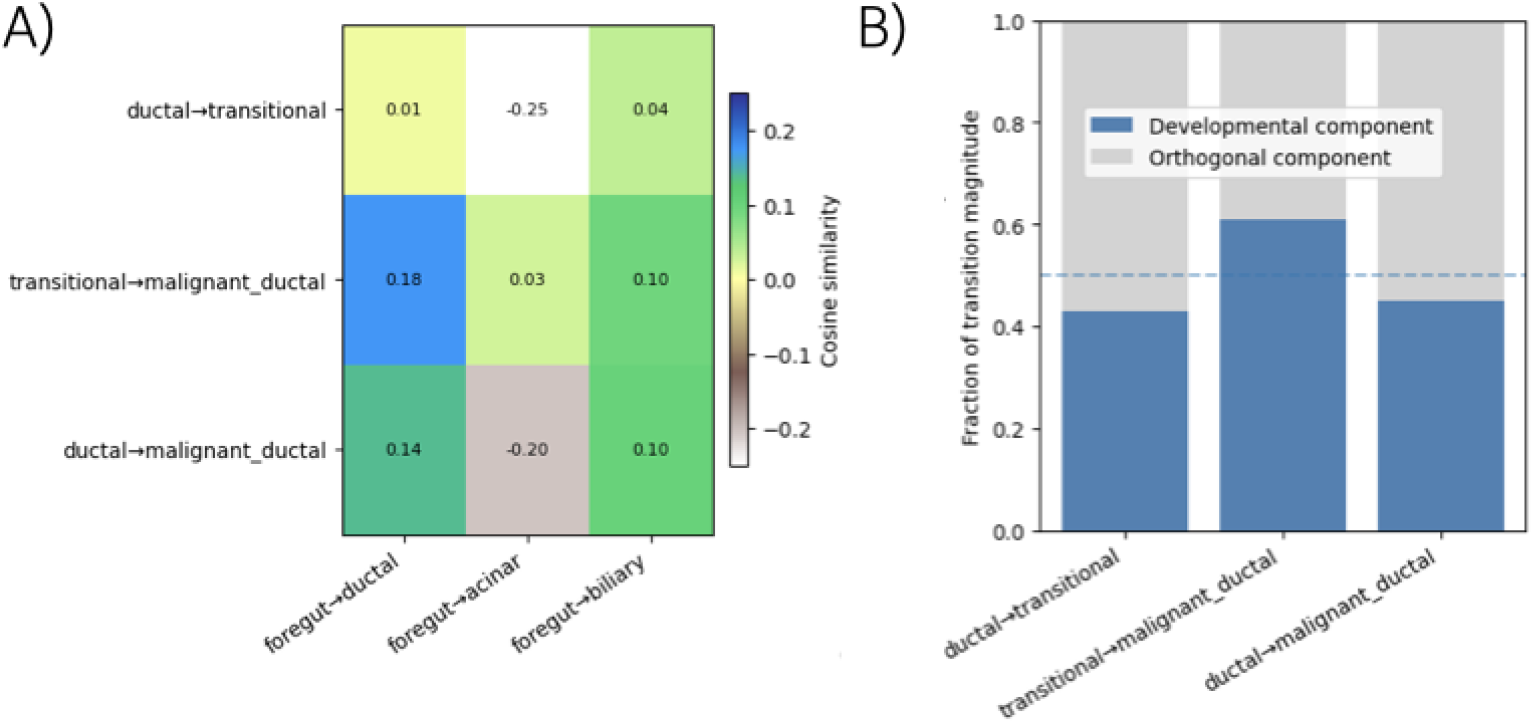
Malignant state differences are weakly aligned with canonical developmental directions and retain substantial components outside the developmental subspace. A) Cosine similarity between malignant centroid-to-centroid displacement vectors and three developmental branch directions. Cosine values remain small across comparisons, indicating weak alignment between malignant state differences and canonical developmental axes. B) Decomposition of the same malignant displacement vectors into components within the developmental PC subspace and outside it. Developmental structure captures a measurable fraction of each displacement, but substantial residual components remain outside the developmental subspace.

Together, these analyses show that malignant-state differences partially reuse developmental structure but do not follow canonical branch directions. The residual component of these differences is therefore not a technical remainder. It represents the component of variation that is not captured by the developmental reference space and is analyzed biologically below.

### Developmental geometry explains malignant state positions but not malignant state variation

The preceding analyses distinguish two related but different questions. The first is positional: where do cancer states lie within a developmental reference frame? The second is directional: do deviations between states follow differentiation trajectories? Developmental geometry helps answer the first question, but not the second.

We first addressed the positional question. If malignant states retain developmental character, their centroids should occupy interpretable locations relative to lineage directions, even if differences between states do not follow those directions. For each state centroid, we computed relative alignment with the developmental axes, normalized within state (Fig. 3A; Supplement). Cancer states occupied distinct positions within this coordinate system. The ductal state retained stronger acinar/secretory-associated positioning, whereas the malignant ductal state showed stronger ductal-associated positioning. Thus, geometry provides a useful coordinate system for describing malignant-state identity, but not a trajectory model of tumor progression.

**Figure 3:**
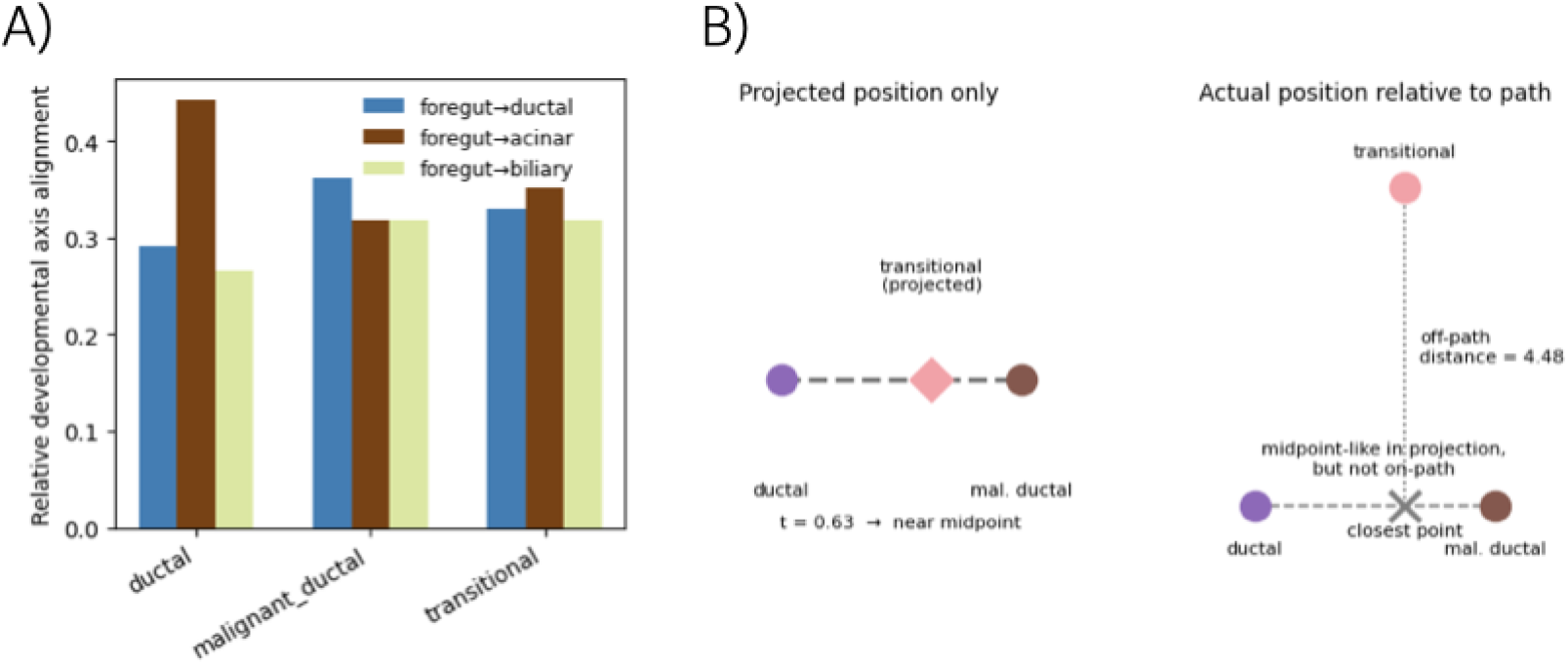
Developmental geometry describes malignant state positions but does not imply developmental trajectory. A) Relative alignment of each malignant-state centroid with three canonical developmental axes: foregut-to-ductal, foregut-to-acinar, and foregut-to-biliary. Alignment scores were normalized within each malignant state, so the plot shows relative developmental-axis preference rather than absolute developmental constraint. This representation summarizes where each malignant state lies within a simplified developmental coordinate system. B) Test of whether the transitional state is a geometric intermediate between ductal and malignant ductal states in developmental PC space. Left, after projection onto the one-dimensional ductal-to-malignant ductal axis, the transitional centroid falls between the two endpoints (*t* = 0.63). Right, in the full developmental PC space, the transitional centroid lies substantially off the endpoint-connecting path. The red × marks the closest point on the path, and the dotted segment indicates the perpendicular deviation (*d*_⊥_ = 4.48). Thus, the transitional state appears intermediate only after one-dimensional projection; it is not collinear with the ductal and malignant ductal states in developmental space.

To interpret these positions biologically, we use gene expression, curated developmental and cancer programs, malignant-state enrichment, and residual cancer-associated axis con-tributions (Supplement, Table S1; Fig. S10). State identity was shaped by both developmental-like positioning and residual cancer-associated programs. The *ductal* state retained acinar/secretory digestive enzyme identity, including expression of Pnlip, Zg16, and Aqp12, and showed strong enrichment along an acinar/secretory residual axis. The *malignant_ductal* state combined ductal/biliary maturation alignment with glycolytic, hypoxic, oxidative-stress, MYC, oxidative-phosphorylation, and stemness/progenitor-like programs. The *transitional* state showed weak canonical developmental alignment but carried neuroendocrine-like markers, including Gip, Fev, and Neurod1, together with modest p53, AP-1/KRAS, ECM-remodeling, unfolded-protein-response, and antigen-presentation activity. These annotations show that the cancer-state phenotype combines lineage-like features with cancer-adaptive programs not captured by canonical developmental branch directions.

The positional signal was also broader than the three canonical lineage axes. The span of the three lineage axes captured a mean fraction of 0.216 of the foregut-relative malignant-cell displacement, whereas the full subspace captured a mean fraction of 0.404 of total malignant-cell magnitude (Fig. 1C; Supplement). Thus, cancer cells are not explained simply by proximity to ductal, acinar, and biliary lineage axes; their developmental component is distributed across broader modes of epithelial variation.

We next tested whether a malignant state that appears intermediate in a simplified projection is also intermediate in the full developmental space. When the transitional centroid was projected onto the one-dimensional ductal-to-malignant ductal endpoint axis, it appeared intermediate, lying 62.7% of the way from ductal to malignant ductal (Fig. 3B, left). However, in the full 75-dimensional space, it was substantially displaced from the endpoint-connecting line, with a perpendicular deviation of 4.479, corresponding to 64.1% of the ductal-to-malignant ductal centroid separation (Fig. 3B, right), and a 45.6^*°*^ angle between the ductal-to-transitional and ductal-to-malignant ductal directions (Supplement). Thus, apparent intermediacy in one-dimensional projection does not imply a simple differentiation trajectory.

Finally, we annotated biological programs associated with malignant-state displacement vectors (Supplement, Fig. S11). For each pairwise displacement, we projected cancer cells onto the corresponding vector and correlated these coordinates with gene expression and curated developmental and cancer-associated programs. The ductal-to-transitional displacement, starting from an acinar/secretory-like malignant state, was associated with loss of acinar/secretory digestive enzyme identity and gain of AP-1/KRAS, p53, inflammatory, senescence/SASP, epithelial, and unfolded-protein-response programs. The transitional-to-malignant ductal displacement was associated with increasing glycolytic rewiring, oxidative stress, hypoxia, EMT/invasion, cell-cycle activity, stemness/progenitor-like identity, and MYC target programs. Across the full ductal-to-malignant ductal displacement, malignant cells lost acinar differentiation and secretory digestive enzyme programs while gaining ductal/foregut epithelial, progenitor-like, metabolic, hypoxic, oxidative-stress, MYC, senescence, and invasive programs. Thus, malignant-state variation reflects cancer-adaptive reprogramming superimposed on developmental-state repositioning, rather than movement along normal differentiation paths.

## Discussion

Our results refine the relationship between normal tissue programs and malignancy by showing that those programs provide an informative reference for malignant epithelial identity but do not fully define the directions along which cancer states diversify. This distinction between positional similarity to normal cells and the directions of pathological divergence follows from comparing the organization of normal epithelial variation with how transformed states separate from one another.

Developmental cells are well described by a relatively low-dimensional reference space, compatible with constrained differentiation. Malignant cells, by contrast, are only partially captured by this space: a substantial fraction of their embedding magnitude lies outside the dominant axes of developmental organization. This residual component is also low-dimensional, biologically *structured*, and informative of cancer-state identity. PDAC therefore extends along transcriptional directions beyond those defining normal epithelial programs.

Malignant PDAC states have been described as co-opting acinar, pancreatic progenitor, endocrine-progenitor, basaloid, and squamoid programs, with basal and squamous-like states proposed to be accessible through broader foregut developmental relationships (Patel et al. 2024). Our results are compatible with this broadened positioning, but indicate that it is not sufficient to explain tumor-driven diversification. The three foregut-to-terminal lineage axes did not exhaust the developmental signal present in malignant cells; a broader developmental PC subspace captured substantially more structure. Thus, the developmental component of malignant character is distributed across the reference space rather than reducible to a small number of canonical lineage directions.

An aberrant cell state may therefore occupy a recognizable developmental neighborhood without being generated by, or progressing along, the corresponding developmental trajectory. This distinction helps interpret basal or squamous-like PDAC states: their relationship to foregut-associated epithelial programs may help explain their position, but it does not by itself establish a trajectory through pancreatic-esophageal progenitor-like states. More generally, these results challenge the assumption that the structure used to position cells also constrains the directions of cellular change.

This view is aligned with work emphasizing that PDAC progression is driven by cellular plasticity, including acinar-to-ductal metaplasia, subtype specification, early dissemination, metastatic-state plasticity, and treatment resistance (Jiménez-Sánchez et al. 2026; Innamorati et al. 2026; Gautam et al. 2026). Our analysis addresses a complementary question: how is this plasticity organized relative to normal epithelial maturation? If malignant plasticity were primarily organized by constraint, one would expect malignant-state differences to align strongly with developmental axes. Instead, developmental coordinates help locate tumor states, whereas the dominant directions of separation involve additional cancer-associated programs. Thus, PDAC plasticity is not simply movement along normal developmental trajectories, but a reconfiguration of epithelial identity through stress-adaptive, invasive, metabolic, inflammatory, and proliferative axes. In this sense, we extend developmental-reuse and -constraint models of cancer cell identity (Tirosh et al. 2024; Neftel et al. 2019; Patel et al. 2024).

The distinction also clarifies the interpretation of apparently intermediate tumor states. In our analysis, the transitional malignant state appears intermediate when reduced to a one-dimensional ductal-to-malignant ductal coordinate, but deviates substantially from the endpoint-connecting path in the full developmental subspace. Apparent continuity in projected space can therefore arise even when states are not true geometric intermediates.

Moreover, the biological interpretation benefited from MLM embeddings that capture transcriptional *context* and coordinated gene programs rather than single markers alone. This motivation is related to broader quantitative efforts to represent cellular states and transitions using manifold learning, optimal transport, RNA velocity, and representation learning (Moon et al. 2019; Schiebinger et al. 2019; Qiao et al. 2021; Burkhardt et al. 2021). However, our goal was not to infer a trajectory directly, but to test whether malignant-state differences align with a normal developmental reference. We therefore evaluated the embedding by biological interpretability: developmental PCs recovered expected epithelial programs, malignant residual axes captured known cancer-associated programs, and both projected and residual components retained information about cancer-state identity.

These conclusions should be interpreted, however, in light of several limitations. The reference space is a linear approximation and may not capture nonlinear aspects of epithelial differentiation. The embedding is learned from jointly modeled developmental and cancer datasets, so aspects of the representation may reflect dataset composition, although the goal of the model is precisely to learn a shared space in which both biological features can be compared. Displacement vectors are inferred from static single-cell measurements and should not be interpreted as directly observed temporal trajectories. In addition, directions outside the reference space are defined relative to the *specific* developmental dataset considered and may include normal biological variation absent from this reference. Finally, although the residual axes are biologically structured, their functional role in tumor progression remains to be experimentally tested. Future work combining this decomposition with lineage tracing, perturbation, and chromatin accessibility measurements could test whether these axes reflect regulatory constraints, adaptive responses to tumor microenvironmental pressure, or both.

More broadly, we provided a framework for comparing disease states with normal biological variation. By decomposing cellular embeddings into components aligned with a reference space and components outside it, one can distinguish shared coordinates of identity from disease-associated directions of variation. In PDAC, developmental variation provides an interpretable basis for positioning malignant states, including those with pancreatic and broader foregut-associated features, but it does not capture the dominant geometry of malignant diversification.

## Methods

### Embedding space and datasets

Cell embeddings were obtained from a pretrained BERT-based MLM (Devlin et al. 2019) trained on large-scale single-cell data. MLM learning was assessed by masked-token reconstruction loss and top-k accuracy, averaged across 10 masking seeds and compared with an architecture-matched random model. We analyzed differentiating epithelial cells from the mouse embryonic and fetal atlas of (Qiu et al. 2024) and malignant epithelial cells from the pancreatic cancer progression cohort of (Burdziak et al. 2023). For each cell, the model-derived CLS embedding was used as its vector representation. Note also that developmental/cancer labels are not used to train the embedding, only after embedding for interpretation.

### Developmental space

Developmental and cancer embeddings were centered using the mean developmental embedding. PC analysis was performed on centered developmental embeddings to define a linear developmental reference subspace. Cancer embeddings were then decomposed into a component projected into this developmental subspace and an orthogonal residual component. The fraction of squared embedding magnitude captured by the developmental projection was used as a *per*-cell developmental projection score. PC of cancer residuals was used to assess whether non-developmental malignant variation was structured.

### Developmental directions

To define biologically interpretable directions, we computed centroids for foregut, ductal, acinar, and biliary developmental populations. Lineage directions were defined as centroid-to-centroid vectors from foregut to each terminal epithelial state. Pairwise angles and bootstrap resampling were used to assess the stability of these directions. Each developmental or malignant cell was assigned a longitudinal coordinate and an orthogonal distance relative to each lineage direction, allowing cells to be positioned within the developmental branch coordinate system.

### Cancer transitions

Cancer-state centroids were computed for annotated cancer states. Transition vectors between malignant states were defined as centroid-to-centroid differences. We quantified their alignment with developmental lineage directions using cosine similarity. We also decomposed each transition vector into components within and outside the developmental PC subspace to distinguish broad developmental-subspace overlap from alignment with specific canonical lineage axes.

### Malignant structure and developmental directions

To test whether cancer-cell structure could be explained by the three lineage axes alone, we compared reconstruction by the full developmental PC subspace with reconstruction by the span of the three foregut-to-terminal lineage vectors, using the Moore–Penrose pseudoinverse for the non-orthogonal axis matrix. Finally, to determine whether transitional malignant states were true geometric intermediates, we projected ductal, transitional, and malignant ductal centroids into devel-opmental space and measured both the scalar position of the transitional centroid along the ductal-to-malignant ductal line and its perpendicular deviation from that line.

Additional biological background, datasets information and analyses, and full mathematical details are provided in the Supplement.

## Acknowledgments

This work was supported by grant PID2023-151289NB-I00 funded by MICIU/AEI/10.13039 /501100011033 and by “ERDF/EU”.

## Code availability

Code available at github.com/juanfpoyatos/OncoDevFormer. This work was done with a single GPU unit. The author used OpenAI’s ChatGPT to assist with text improvement and code debugging. The author reviewed and edited all content and is fully responsible for the final manuscript.

