## Supplement for "Development positions malignant cellular states but does not explain their diversification"

The following supplement provides the biological background and characterization required for this study, together with supplementary information on the data, the relational approach with the use of language models, and the mathematical analyses.

### Biological background and reference framework

The analyses in the main text compare malignant pancreatic epithelial states with developmental epithelial variation. We now summarize the biological background motivating the developmental references used and the interpretation of residual programs.

#### Pancreatic epithelial development and the choice of developmental axes

The pancreas develops from foregut endoderm. In mice, pancreatic specification begins around embryonic day 8.5, followed by formation of dorsal and ventral pancreatic buds. These epithelial primordia generate the major pancreatic compartments, including *acinar* cells, *ductal* cells, and endocrine *islet* cells. Acinar cells form the enzyme-producing exocrine compartment, ductal cells form the branched ductal network, and endocrine cells generate hormone-producing islets (Gittes 2009; Pan et al. 2011; Shih et al. 2013; Stanger et al. 2013). During development, multipotent pancreatic progenitors become spatially and transcriptionally organized into tip and trunk domains, with tip progenitors biased toward acinar differentiation and trunk progenitors contributing to ductal and endocrine lineages (Zhou et al. 2007; Schaffer et al. 2010; Jennings et al. 2013; Ma et al. 2023). These developmental relationships motivate the use of *foregut-to-acinar* and *foregut-to-ductal* directions as biologically interpretable pancreatic epithelial differentiation axes.

A biliary reference is also biologically relevant in this setting. The ventral pancreatic bud develops near the liver and biliary domain and is anatomically associated with the common bile duct. Pancreatic, hepatic, and biliary epithelia therefore arise from neighboring foregut territories shaped by related patterning signals (Gittes 2009; Zaret et al. 2008; Pan et al. 2011; Jennings et al. 2013). In this study, the *foregut-to-biliary* direction is not used to imply that pancreatic ductal adenocarcinoma (PDAC) follows a biliary developmental trajectory. Rather, it provides an adjacent foregut-derived pancreatobiliary epithelial reference that helps distinguish ductal, acinar, and biliary-like components of malignant identity.

This choice of axes is intentionally simplified. The foregut-to-ductal, foregut-to-acinar, and foregut-to-biliary directions do not exhaust the complexity of pancreatic and pancreatobiliary development. They summarize three biologically interpretable directions within a broader developmental PC-coordinate system. As shown in the main text and Supplementary analyses, these axes are used to ask whether cancer-state differences align with canonical developmental directions, whereas the full developmental PC subspace is used to ask whether malignant variation is captured by *broader* developmental structure.

#### PDAC progression, acinar-to-ductal plasticity, and developmental resemblance

PDAC is commonly associated with ductal morphology and precursor lesions such as pancreatic intraepithelial neoplasia (PanIN). However, experimental and lineage-tracing studies have shown that acinar cells can serve as an important cellular origin for pancreatic tumorigenesis (Hruban et al. 2000; Maitra et al. 2008; Kleeff et al. 2016; Kopp et al. 2012; Storz 2017). In response to injury, inflammation, and oncogenic KRAS signaling, differentiated acinar cells can undergo acinar-to-ductal metaplasia (ADM), a normally reversible regenerative process in which acinar cells lose mature digestive-enzyme identity and acquire duct-like or progenitor-like features (Reichert et al. 2011; Storz 2017; Marstrand-Daucé et al. 2023). When combined with persistent oncogenic signaling, chronic inflammation, or tumor-suppressor loss, this plasticity can become fixed and promote PanIN formation and progression toward PDAC (Morris et al. 2010; Shi et al. 2019; Kopp et al. 2012).

Recent work has emphasized that PDAC progression cannot be understood as a simple linear accumulation of driver mutations. Instead, early neoplastic progression involves phenotypic plasticity, inflammatory and stromal cues, ADM, epithelial–mesenchymal transition (EMT)-like programs, squamous or basal-like transdifferentiation, and early dissemination from precursor or early malignant lesions (Bärthel et al. 2023; X. Zhang et al. 2025; Innamorati et al. 2026). This perspective reinforces the need to distinguish developmental resemblance from cancer-state dynamics. A cell state may display ductal, acinar, progenitor-like, or pancreatobiliary features while also acquiring stress-adaptive, invasive, metabolic, or mesenchymal programs that are not equivalent to normal developmental trajectories.

This biology motivates the comparison between malignant states and developmental epithelial references. PDAC cells may retain or reacquire lineage-associated programs, including acinar/secretory, ductal, foregut-like, progenitor-like, or pancreatobiliary features (Bailey et al. 2016; Chan-Seng-Yue et al. 2020; Burdziak et al. 2023). However, developmental resemblance alone does not establish that cancer states move along normal developmental trajectories. The analyses in the main text therefore distinguish developmental *position*, meaning where malignant states lie relative to developmental epithelial references, from state *direction*, meaning how cancer states differ from one another.

#### Plasticity, inflammation, and the PDAC tumor microenvironment

PDAC is characterized by extensive epithelial plasticity and by a dense, interactive tumor microenvironment. Cells interact with cancer-associated fibroblasts, macrophages, neutrophils, T cells, endothelial cells, extracellular matrix, and other stromal components. These interactions occur through cytokines, growth factors, extracellular matrix remodeling, immune-suppressive signaling, metabolic constraints, and hypoxic stress (Feig et al. 2012; Hessmann et al. 2020; Steele et al. 2020; Truong et al. 2021). The resulting microenvironment can reinforce malignant epithelial plasticity, promote partial EMT, support invasive and stem-like states, and contribute to therapeutic resistance (Öhlund et al. 2017; Biffi et al. 2019; Elyada et al. 2019; Marine et al. 2020; Raghavan et al. 2021; T. Zhang et al. 2022).

Inflammation is particularly important in PDAC initiation and progression. Injury-associated and inflammatory signaling can promote ADM, cooperate with oncogenic KRAS, and accelerate progression toward PanIN and invasive carcinoma (Storz 2017; Shi et al. 2019; Marstrand-Daucé et al. 2023; Burdziak et al. 2023). Microenvironmental remodeling, including desmoplasia, extracellular matrix deposition, altered tissue mechanics, and hypoxia, can further shape cancer cell states and promote invasive behavior (Whatcott et al. 2015; Hessmann et al. 2020; Panciera et al. 2020; Masugi 2022). These observations provide biological context for the residual axes identified in the main text, which include AP-1/KRAS signaling, inflammatory and antigen-presentation activity, hypoxia, glycolytic rewiring, oxidative stress, EMT/extra-cellular-matrix (ECM) remodeling, MYC-associated proliferation, and unfolded protein response programs.

The same framework also highlights the importance of tumor-microenvironmental and metastatic-site control of plastic states. Inflammatory signaling, ECM remodeling, desmoplasia, hypoxia, metabolic stress, immune interactions, and metastatic-site cues can promote EMT-like and squamous/basal-like programs, early dissemination, dormancy, treatment resistance, recurrence, and transcriptomic plasticity in metastatic disease (Rhim et al. 2012; Aiello et al. 2018; J. Yang et al. 2023; Pastushenko et al. 2019; Jiménez-Sánchez et al. 2026; Innamorati et al. 2026). These processes are not simply developmental lineage programs. They represent *cancer-adaptive responses* that may be superimposed on lineage-associated identity. Thus, residual variation outside the developmental reference space is expected to contain structured cancer-associated biology rather than only technical noise.

EMT and partial EMT-like programs, metabolic rewiring, and stress adaptation are well-established components of cancer plasticity and PDAC progression (Ying et al. 2012; Son et al. 2013; Nieto et al. 2016; Pastushenko et al. 2019; J. Yang et al. 2023). This is important for interpreting the residual component in the main text. The residual should not be read as 'non-biological' simply because it lies outside the developmental reference space. Rather, it captures malignant programs that are plausibly induced by oncogenic signaling, inflammation, metabolic stress, tissue mechanics, immune interaction, and stromal remodeling.

#### Implications for interpreting the geometric analysis

This developmental and disease context motivates the geometric tests in the main text. Foregut-to-acinar and foregut-to-ductal directions summarize major pancreatic epithelial differentiation axes, whereas the foregut-to-biliary direction provides an adjacent pancreatobiliary epithelial reference. These axes provide a simplified but biologically interpretable coordinate system for comparing malignant epithelial states with normal developmental variation.

If PDAC plasticity were primarily a replay of normal development, cancer-state differences would be expected to align with these developmental directions or to be largely contained within the developmental reference space. Conversely, if plasticity combines developmental identity with cancer-adaptive reprogramming, malignant states could still occupy interpretable developmental positions while differing along additional axes associated with KRAS signaling, inflammation, metabolic stress, hypoxia, EMT, and tumor-microenvironment interaction.

The main text evaluates these alternatives by separating two questions. The first is *positional*: where do cancer states lie relative to developmental epithelial references? The second is *directional*: do the differences between states follow developmental lineage directions? This distinction is essential because developmental resemblance does not by itself imply developmental trajectory. A state can be ductal-like, acinar-like, biliary-like, or progenitor-like in position while still differing from other malignant states along cancer-adaptive directions. In this study, the developmental reference space provides an interpretable coordinate system for identity, whereas the residual and state-displacement analyses test whether malignant diversification follows, or departs from, normal developmental geometry.

### Datasets

#### Endodermal epithelial reference atlas

**Data source.** We used the mouse embryonic scRNA-seq compendium from Qiu et al. 2024, which spans 11.4 million cells from embryonic (E) day 8 to birth (P0) and integrates with previously published datasets covering E0-E8.5. These authors constructed a global tree of 283 cell-type nodes across 14 developmental subsystems, representing all major lineages of the mouse embryo. Our analyses draw specifically from the organogenesis and fetal epithelial subsystems within this resource.

**Rationale for a focused epithelial/endodermal subset.** Our objective was to train a masked-language model (MLM) on the transcriptional 'grammar' of endoderm-derived epithelial tissues related to pancreatic identity. Because Qiu *et al.*'s global reference includes many non-endodermal and non-epithelial tissues (mesoderm, neural crest, sensory epithelium, immune lineages), using the full dataset would introduce orthogonal expression programs that dilute the foregut- and gut-relevant signal. We therefore extracted a curated subset of epithelial and endodermal populations, chosen based on developmental lineage, organ identity, and relevance for pancreatic differentiation.

Critically, the available E8-P0 portion of the dataset contains mature and fetal epithelial states but does not include early foregut or pancreatic progenitors (e.g., definitive endoderm, primitive gut tube, hepatic/pancreatic buds). Our atlas therefore represents a mature-to-fetal foregut/gut epithelial manifold, not a full zygote-to-epithelium developmental trajectory.

**Selection of cell types.** We filtered the 'df_cell.csv' metadata (Qiu et al. 2024) and retained only cell types belonging to foregut- or gut-derived epithelia, including pancreas, hepatobiliary system, lung/airway, and intestinal epithelium, plus pancreatic islets. The whitelist included:

- Pancreas: pancreatic acinar cells; foregut epithelial cells (ductal proxy); pancreatic islets.
- Hepatobiliary: hepatocytes; biliary epithelial cells.
- Gut: Gut; Midgut/Hindgut epithelial cells; intestinal goblet and enteroendocrine cells.
- Lung: lung progenitor cells; alveolar Type I/II cells; airway club and goblet cells; $Eln^{+}$ lung epithelial cells.
- Foregut-associated endodermal epithelia: thyroid, parathyroid, and thymic epithelial cells.

Ectodermal epithelia (keratinocytes, dental/otic/olfactory epithelium), mesoderm-derived epithelia (kidney, ureteric bud, bladder), extraembryonic endoderm, immune lineages, endothelium, and all mesenchymal populations were excluded.

**Annotation of coarse categories.** Each retained cell type was mapped into one of the following coarse epithelial categories:

- Pancreatic: acinar, ductal-proxy, islet.
- Hepatobiliary: hepatocyte, biliary epithelium.
- Pulmonary: lung airway epithelium (alveolar and conducting); lung progenitor.
- Intestinal: proximal/mixed gut epithelium; distal gut epithelium (midgut/hindgut).
- Foregut-other: thyroid, parathyroid, thymic epithelium.

These categories were defined using a combination of exact cell-type matches and substring-based fallback rules. A final set of allowed categories ('ALLOW_ENDO_CATEGORIES') enforced exclusion of all off-trajectory tissues.

**Merging with expression data.** For each raw '.h5ad' file provided, cell IDs were normalized by stripping run-specific suffixes to create a stable 'cell-id-base'. Metadata rows were then merged onto AnnData objects using this key. Only cells whose 'cell_id_base' matched curated metadata entries were retained.

**Developmental time annotation.** Developmental day information (E and P stages) was harmonized by parsing expressions of the form 'E18.5' or 'P3.5'. Postnatal days were offset by +20 to place them after embryonic stages on a continuous axis ('dev_day').

**Pancreas-axis label.** We defined a pancreas-specific state ('dev_state_pancreas') with three levels. This label was overlaid onto all pancreatic cells: 1/acinar_or_tip, 2/ductal, and 3/non_epithelial_endocrine (islets)

**Union label for downstream modeling.** A combined label ('dev_state_union') was created that uses the pancreatic state when available, or else falls back to the coarse epithelial category. This ensures that each cell is assigned one biologically interpretable category for visualization and MLM training.

**Downsampling for class balance.** Because hepatocytes and gut epithelium dominate the retained dataset, we subsampled each 'dev_state_union' category to at most 4,000 cells using stratified random sampling. This produced a balanced atlas ('dev_endoderm_balanced.h5ad') with comparable representation across epithelial classes for MLM training.

**The mouse developmental epithelial manifold.** Our atlas provides a structured representation of mature-to-fetal epithelial programs that define the physiological neighborhood within which pancreatic cells normally exist, and which PDAC cells are known to aberrantly explore. To assess its structure, we performed principal component (PC) analysis and uniform manifold approximation and projection (UMAP) on the cells belonging to the selected epithelial categories. The resulting manifold exhibited clear, biologically interpretable organization (Fig. S1). Hepatocytes formed a compact block adjacent to biliary epithelium, reflecting their shared hepatobiliary lineage. Proximal/mixed gut epithelium and distal gut epithelium separated cleanly along an intestinal axis. Lung progenitors and airway/alveolar epithelium formed a continuous arc spanning differentiation within the respiratory lineage. Pancreatic acinar and ductal-proxy populations appeared as distinct islands near hepatobiliary and proximal gut tissues, consistent with their foregut origin, while pancreatic islets formed a separate endocrine outgroup. Thyroid, parathyroid, and thymic epithelial cells grouped together as a foregut-associated cluster. The lack of admixture between unrelated tissues and the continuity within organ-specific arcs confirm that the atlas captures the expected epithelial relationships across foregut, midgut, and hindgut territories.

Overall, this epithelial manifold captures the major transcriptional axes that PDAC is known to traverse, including acinar-ductal transitions, biliary and hepatocyte-like programs, gastric/intestinal shifts, and occasional lung-like or endocrine-like states. By training a MLM (see below) on this structured and lineage-constrained reference, we establish a semantic space in which PDAC states can be interpreted not relative to tumor-internal clusters, but relative to the broader epithelial grammar encoded during mammalian development.

#### Pancreatic ductal adenocarcinoma dataset

**Data source.** We considered the progression cohort in Burdziak et al. 2023, a single-cell analyses on normal, inflamed, premalignant, and malignant tissues in a genetically engineered mouse model –that develop tumors spontaneously within the organism, mimicking natural disease processes– of PDAC. This dataset spans 28,131 cells.

**Epithelial analysis.** To evaluate whether this progression cohort (Burdziak et al. 2023) primarily contained epithelial tumor cells, we first defined a canonical epithelial marker panel (Epcam, Krt8, Krt18, Krt19, Cdh1) –specifically, EpCAM and keratins, like Krt8, Krt18, and Krt19, are often associated with epithelial tissues, while Cdh1, E-cadherin, is important for cell adhesion in epithelial cells.

We then computed a *per*-cell epithelial score ('epi_score') using Scanpy (Wolf et al. 2018). Across all cells, epi_score values were broadly distributed (mean 16.1, IQR 4.0–24.9). We then considered a mild epithelial gate based on the 30th percentile of this score (threshold 6.315), which would retain the top 70% of cells (19,692 cells) and discard 8,439 low-scoring cells. To verify whether this low-epi fraction corresponded to non-epithelial contamination, we examined the expression of canonical immune (Ptprc, Cd3d, Cd68, Lyz), fibroblast (Col1a1, Col1a2, Pdgfra, Acta2), endothelial (Pecam1, Kdr, Vwf), and EMT/mesenchymal (Vim, Zeb1, Snai1, Snai2) markers within the low-epi subset using violin plots stratified by condition.

This analysis did not reveal a dominant immune, fibroblast, or endothelial signature confined to the low-epi fraction, but instead suggested a mixture of lower-coverage epithelial cells and cells with partial or EMT-like epithelial programs. Given the prior epithelial enrichment of the dataset and our goal of modeling the full spectrum of malignant epithelial plasticity, we therefore did not apply an epi_score-based hard filter. Instead, we retained all 28,131 cells for model training and used epithelial scores and lineage markers as diagnostic annotations in downstream embeddings, with the option to remove clearly non-malignant clusters at the cluster level if required. Moreover, epithelial identity varied continuously across the cancer manifold and did not define a separable cell population; therefore epithelial scores were used as diagnostic annotations rather than hard inclusion criteria.

**The PDAC manifold.** To characterize the global transcriptional landscape of pancreatic tumorigenesis in the progression cohort, we constructed a UMAP representation of the full epithelial compartment, including both lineage-traced epithelial cells (mKate2$^+$) and epithelial-enriched samples (EPI). Unlike the lineage-restricted visualization shown in the original study, which focused primarily on mKate2$^+$ cells and emphasized heterogeneity within malignant programs, this embedding captures the entire epithelial trajectory spanning normal pancreas, injury-induced metaplasia, preneoplastic lesions, primary tumors, and metastases (Fig. S2).

Cells organize along a continuous progression from normal epithelium (N1) to regenerating pancreas (N2), followed by early KRAS-driven metaplasia and preneoplasia (K1-K2), which form a broad transitional compartment linking normal ductal states to neoplastic programs. Advanced lesions diverge into distinct transcriptional territories corresponding to invasive PDAC (K5) and metastatic disease (K6). Coloring the embedding by inferred transcriptional state further reveals a hierarchy of epithelial phenotypes, with canonical ductal cells occupying the normal and regenerative regions, transitional states dominating the preneoplastic landscape, and malignant ductal programs characterizing invasive and metastatic tumors. The resulting manifold therefore captures both the temporal ordering of tumor progression and the gradual remodeling of epithelial identity during pancreatic tumorigenesis.

Note also how normal pancreas cells (N1) display relatively low epithelial scores, reflecting the mixture of acinar, endocrine, and ductal cell types, only some of which strongly express classical epithelial markers. During early tumorigenesis (N2–K2), epithelial scores increase as acinar cells undergo acinar-to-ductal metaplasia (ADM) and activate ductal epithelial programs. The highest epithelial scores appear in intermediate preneoplastic states (K2–K4), where cells adopt strong ductal epithelial identities characteristic of PanIN lesions. In contrast, invasive PDAC (K5) shows a modest reduction in epithelial scores, consistent with partial EMT programs associated with malignancy. Metastatic cells (K6) display intermediate epithelial scores, suggesting hybrid epithelial-mesenchymal phenotypes. Overall, the epi_score panel highlights a progressive remodeling of epithelial identity along the trajectory from normal pancreas through preneoplastic lesions to invasive and metastatic PDAC.

#### Data processing and normalization

**Construction of a shared gene set.** The developmental reference atlas from Qiu et al. 2024 was processed using the mouse reference genome mm10 (GRCm38) and GENCODE VM12 gene annotations, yielding count matrices with Ensembl gene identifiers in the feature dimension. In contrast, the PDAC single-cell dataset from Burdziak et al. 2023 were generated on the older mm9 (NCBI37) genome assembly using a RefSeq-/RIKEN-based gene set, and the released count matrices use heterogeneous gene symbols (including historical RIKEN clone IDs, predicted mouse gene loci, GenBank accessions and hybrid human-style symbols). To enable joint analysis of these data, we reconciled both datasets onto a common, *modern* annotation framework based on GRCm39 and Ensembl release 111.

We first parsed the Ensembl GTF 'Mus_musculus.GRCm39.111.gtf' to build maps from Ensembl gene IDs to gene symbols and from gene symbols to Ensembl IDs. Only gene features were retained, and Ensembl version suffixes (e.g., ENSMUSG00000025945.1) were stripped to yield stable gene IDs. For the developmental atlas, we replaced the original names with version-stripped Ensembl IDs and attached the corresponding Ensembl gene symbols. For the PDAC dataset, we mapped each symbol to a GRCm39 Ensembl ID in a stepwise fashion. First, we performed a case-insensitive lookup against the GRCm39 gene_name field using the symbol to ensemble dictionary constructed from the GTF. Symbols that were not found at this step typically correspond to historical identifiers that no longer exist as primary gene names in GRCm39. To recover a subset of these, we optionally queried Ensembl's REST API and MyGene.info for each unmapped symbol, retrieving Ensembl gene IDs when a unique, high-confidence match was available; these external matches were incorporated as an additional manual alias map. Symbols that remained unmapped after the GTF-based and external alias steps were excluded from all joint developmental–PDAC analyses but retained in PDAC-only analyses. When multiple PDAC symbols mapped to the same Ensembl ID (for example, different aliases of the same gene), we collapsed the corresponding columns by summing their counts.

We then defined the shared gene universe as the intersection of Ensembl IDs present in both the developmental and PDAC datasets, and restricted each AnnData object to this common set. This produced two aligned expression matrices, 'A_dev_aligned' and 'A_cancer_aligned', with identical Ensembl feature sets and harmonized gene symbols. Overall, starting from 24,552 developmental genes and 16,826 PDAC symbols, this procedure yielded a shared set of 15,376 genes used for all developmental-PDAC comparisons.

**Unification of expression datasets.** One cannot meaningfully compare raw counts between the developmental and cancer datasets because raw counts between the two datasets are not comparable in any biologically meaningful way: 1/they come from different experiments, sequencing depths, capture chemistries, and UMI efficiencies, 2/developmental raw counts have a different global distribution than cancer raw counts (different sparsity, different dynamic range), 3/gene A having '50 counts' in dev and

'50 counts' in cancer is not the same quantity; the total molecule capture per cell can differ by 3-10×.

To unify the datasets, we first apply library-size normalization to each dataset independently (every cell ends up with total 'counts' ≈ 10,000; this is standard CPM-like normalization) and then considered the *developmental data* as the *reference baseline*. We obtain this baseline by computing, for each gene $g$, the nonzero median across developmental cells, median_nonzero($g$); using the developmental dataset after its own internal normalization.

Then, in a given developmental cell $d$, we compute: dev_rel($g$,$d$) = norm_expr($g$,$d$)/median_nonzero($g$), and in a given cancer cell $c$, we compute: cancer_rel($g$,$c$) = norm_expr($g$,$c$)/median_nonzero($g$). Here norm_expr($g$,$d$) and norm_expr($g$,$c$) represent the already library-normalized developmental or cancer expression value, respectively, not the raw count. This expresses each cancer gene's abundance as how many times the typical developmental level a given gene is showing. In other words, it converts cancer expression into relative-deviation coordinates that sit on the developmental manifold learned by the model. This keeps the MLM grounded in the same numerical scale across datasets and makes gene-level deviations interpretable.

Note also that using raw counts in the numerator would turn the developmental ratio into a batch-readout, not a biological deviation; the MLM would learn dataset-specific noise, not developmental grammar. By using normalized expression in the numerator and developmental nonzero medians in the denominator, we ensure that the quantity being compared is on the same approximate scale, controlled for library depth, interpretable as a *fold-shift* relative to typical developmental abundance, and not confounded by experiment-specific artifacts (Theodoris et al. 2023; Poyatos 2025). This is the only way to make the resulting 'grammar deviations' reflect biology, not technical drift.

**Validation of the development-relative normalization.** Because each gene was normalized by its nonzero median expression across developmental cells, the median development-relative (dev_rel) value is expected to equal one by construction. Therefore, the distribution of *per*-gene medians is not an informative diagnostic of normalization quality. Instead, we evaluated the dispersion of normalized developmental values around this fixed baseline using the median absolute deviation (MAD) from one. Across genes with nonzero developmental baselines, MAD values were low, indicating that developmental expression values remained tightly concentrated around their gene-specific reference levels after normalization (Fig. S3A).

We also examined the global distribution of $\log_2$-transformed development-relative expression values across all genes and developmental cells. Since a development-relative value of one corresponds to $\log_2(1) = 0$, a well-behaved normalization should produce a distribution centered near zero. Consistent with this expectation, the distribution was centered near zero and showed a balanced spread without pronounced long tails or distortions (Fig. S3B). These results indicate that the development-relative transformation behaves as expected and does not introduce systematic inflation or compression of gene-level expression values.

**Relative-expression normalization preserves developmental marker specificity while revealing selective malignant deviations.** We next used marker-level controls to evaluate whether relative-expression normalization preserved recognizable developmental structure and produced interpretable malignant deviations from that structure. In the development-relative representation, each gene is scaled by its nonzero median expression across developmental cells, such that a value of one corresponds to the gene-specific developmental baseline. Therefore, this transformation should not make all genes appear uniformly expressed across tissues; instead, broadly expressed epithelial markers should remain broadly distributed, whereas lineage-restricted markers should retain their expected specificity.

Consistent with this expectation, pan-epithelial and ductal-associated markers showed comparable development-relative expression across multiple epithelial endodermal categories. Krt19 and Sox9 were broadly detected across epithelial populations, with values distributed around the developmental baseline rather than being dominated by a single tissue compartment (Fig. S4A). In contrast, lineage-restricted genes retained their expected developmental specificity. The endocrine marker Gcg was enriched primarily in pancreas-islet cells, whereas the hepatic marker Alb was strongly concentrated in liver-hepatocyte cells. Thus, development-relative scaling places genes on a common relative-expression scale without erasing known developmental marker structure.

We then asked how malignant epithelial states deviate from this developmental reference. In the cancer-relative representation, Krt19 was elevated across PDAC states, with particularly strong expression in malignant_ductal and transitional cells, indicating amplification of epithelial/ductal programs beyond the developmental baseline (Fig. S4B). Sox9 showed more moderate but recurrent elevation, consistent with reinforcement of ductal-progenitor-associated identity. By contrast, Gcg remained near zero across cancer states, indicating absence of endocrine-lineage activation, and Alb remained below the developmental baseline, arguing against inappropriate activation of hepatic identity.

Together, these marker-level controls show that relative-expression normalization preserves expected developmental marker specificity while revealing selective PDAC-associated deviations. Cells do not show a nonspecific upward shift across unrelated lineage programs; instead, they preferentially amplify epithelial/ductal-associated expression while maintaining suppression of endocrine and hepatic identities.

**Cross-dataset scaling at the gene level.** Normalized count values (CPM) are not directly comparable across datasets. To demonstrate this explicitly, we computed *per*-gene nonzero median CPM values separately for developmental and PDAC cells and examined their ratios across genes. This analysis revealed a strong global shift, indicating that CPM space does not provide a shared numerical coordinate system for cross-dataset comparison (Fig. S5A).

To establish a biologically meaningful reference frame, we expressed both developmental and cancer expression values relative to a gene-specific developmental baseline as described above. This transformation anchors expression values to the typical developmental `ON` level of each gene and removes global platform-dependent scaling differences. Importantly, dev-relative scaling does not enforce similarity between cancer and development. Instead, it allows deviations to be interpreted relative to a shared developmental grammar.

When summarizing dev-relative expression across all genes, PDAC shows a strong left-shift, reflecting that malignant cells utilize only a *restricted* subset of developmental programs. However, restricting analysis to genes robustly active in PDAC yields near-centered dev-relative values (Fig. S5B), indicating that the numerical scale is aligned for genes participating in the shared epithelial program. Together, these analyses separate technical cross-dataset mismatch from biologically structured deviation, enabling downstream modeling to focus on cancer-specific distortions of developmental gene programs rather than platform artifacts.

Even after *per*-cell library normalization, housekeeping genes such as Actb and Gapdh were 4-7$\times$ more highly expressed in PDAC than in the developmental atlas, demonstrating substantial differences in capture efficiency between datasets. Meanwhile, tissue-specific genes from non-pancreatic lineages (e.g., Tg, Sftpc, Pga5) were strongly expressed in the multi-organ developmental reference but absent in PDAC, producing large negative CPM shifts unrelated to biology. These two effects –platform bias and tissue-composition bias– produce global distortions that overwhelm genuine biological differences.

The dev-relative scaling removes these distortions by expressing each gene in units of its developmental ON-level, ensuring that downstream token selection and MLM embeddings reflect true biological deviations rather than chemistry or lineage composition.

### Masked language modeling

To model transcriptional dependencies across developmental and malignant epithelial states, we implemented a MLM framework adapted from BERT (Devlin et al. 2019). The key idea was to represent each single cell as a discrete sequence of genes ranked by expression relative to a developmental reference, and then train a transformer to predict masked genes from the remaining cellular context (F. Yang et al. 2022; Theodoris et al. 2023; Cui et al. 2024; Szałata et al. 2024; Poyatos 2025).

A shared gene vocabulary was defined across developmental and cancer datasets after confirming identical aligned gene ordering. Special token IDs were reserved for padding (`PAD`), start-of-sequence (`CLS`, see below), end-of-sequence (`SEP`), and masked positions (`MASK`). In addition, two domain tokens were introduced to indicate whether a sequence originated from developmental or cancer data.

For each cell, the input expression profile consisted of the previously discussed *development-relative* gene values, approximating normalized expression divided by the median developmental expression for the same gene. These values were transformed using $\log_{1p}$ after clipping negative values to zero. Genes with undefined developmental baseline were suppressed by setting their score to zero. The top 128 genes with the highest transformed development-relative values were selected per cell and ordered from highest to lowest score. Each cell was then encoded as a fixed-length token sequence of the form [`DOMAIN`] [`CLS`] $g_1 g_2 \ldots\ldots g_{128}$ [`SEP`], where $g_1 \ldots g_{128}$ denote the selected genes in descending rank order. Sequences were padded where necessary, and an attention mask was generated to distinguish real tokens from padding.

Developmental and cancer sequences were pooled into a joint training corpus. A BERT MLM was trained to reconstruct masked gene tokens from their surrounding context. During batch preparation, 15% of gene tokens were randomly selected for masking, excluding padding, special tokens, and domain tokens. Following the standard MLM procedure, 80% of selected tokens were replaced by `MASK`, 10% by a random gene token, and 10% left unchanged, while the loss was computed only on selected positions.

The transformer architecture used hidden size 256, 6 self-attention layers, 8 attention heads, and intermediate feed-forward dimension 512. Training was performed for 10 epochs using AdamW optimization with learning rate $5 \times 10^{-4}$, weight decay 0.01, linear warmup over the first 6% of training steps, linear learning-rate decay thereafter, and gradient clipping at 1.0. Mixed-precision training (fp16) was used to fit the model on an 8 GB GPU. A *per*-device batch size of 8 and gradient accumulation of 4 steps yielded an effective batch size of 32.

We evaluated masked-token reconstruction after training using the same masking procedure applied during training: 15% of non-special tokens were selected, of which 80% were replaced by MASK, 10% by a random gene token, and 10% left unchanged; loss and accuracy were computed only at selected positions. We averaged performance across 10 independent masking seeds and compared with an architecture-matched randomly initialized model. The trained model showed markedly *lower* MLM loss than the random model (4.637 $\pm$ 0.0012 *vs.* 9.690 $\pm$ 0.0002) and substantially *higher* masked-token prediction accuracy: top-1, 17.84% $\pm$ 0.02% *vs.* 0.0067% $\pm$ 0.0010%; top-5, 29.15% $\pm$ 0.03% *vs.* 0.0265% $\pm$ 0.0009%; and top-10, 36.02% $\pm$ 0.04% *vs.* 0.0632% $\pm$ 0.0015%. Thus, training learned nontrivial statistical structure in the tokenized expression corpus, while biological structure in the resulting representations was assessed separately in the downstream analyses.

This framework allowed the model to learn higher-order gene co-occurrence structure across developmental and cancer cell states, conditioned on domain identity, and to capture transcriptional programs that define shared or domain-specific cellular contexts; see Supplement in (Poyatos 2025) for a brief introduction to MLMs.

**CLS embeddings interpretation.** The [`CLS`] ('classification') token is a special class of token used in machine learning models, particularly those based on the Transformer architecture (Devlin et al. 2019).

It is a token that represents the entire input sentence and is placed at the beginning of the input. In our case, CLS embeddings are vectorial encodings of individual cells derived from a pretrained BERT-based model, where each vector summarizes the gene expression profile of a cell in the learned feature space.

### The developmental subspace

To what extent does the geometry of malignant cell states lie within the manifold defined by normal developmental differentiation? If cancer primarily reuses developmental programs, embeddings should lie largely within the developmental subspace. Conversely, if malignant states introduce novel regulatory programs, a substantial fraction of variation should lie outside that subspace. This section provides the quantitative support for Fig. 1, main text.

**Quantifying the subspace.** Let $X_{\text{dev}} \in \mathbb{R}^{n_d \times p}$ denote the developmental dataset containing $n_d$ cells and $p$ CLS-based embedding dimensions (we considered $p = 256$), and let $X_{\text{can}} \in \mathbb{R}^{n_c \times p}$ denote the corresponding cancer embedding dataset with $n_c$ cells. We first centered both datasets using the *developmental* mean

$$\mu = \frac{1}{n_d} \sum_{i=1}^{n_d} X_{\text{dev},i},$$

with $X_{\text{dev},i}$ being each cell $i$ embedding. The centered matrices are therefore $\tilde{X}_{\text{dev}} = X_{\text{dev}} - \mu$, and $\tilde{X}_{\text{can}} = X_{\text{can}} - \mu$. We performed PC analysis on the developmental data $\tilde{X}_{\text{dev}}$ (Jolliffe 2002). Let

$$B = [\mathbf{b}_1, \dots, \mathbf{b}_k] \in \mathbb{R}^{p \times n_{\text{PC}}}$$

denote the matrix containing the first $n_{\text{PC}}$ developmental PCs. These vectors define a $n_{\text{PC}}$-dimensional *linear* subspace $\mathcal{D}$ that captures the dominant axes of transcriptional variation in development. In our case, $\mathcal{D}$ exhibited strong low-dimensional structure, with only the first 75 PCs capturing approximately 98% of developmental variance (variance within developmental epithelial CLS embeddings, not raw expression). This structure indicates that developmental states lie on a coherent low-dimensional manifold (see main text).

**Cancer states on the developmental subspace.** We decomposed each cancer CLS-based embedding vector into a component that lies within the developmental transcriptional subspace and an orthogonal residual component capturing malignant-specific variation. In matrix form, the first projection of cancer cells onto $\mathcal{D}$ is given as $X_{\text{can}}^{\text{dev}} = \tilde{X}_{\text{can}} B$, and the reconstructed developmental component as $X_{\text{can}}^{\text{recon}} = X_{\text{can}}^{\text{dev}} B^{\top}$. This represents the part of the cancer transcriptome that can be explained by developmental transcriptional programs.

When malignant embeddings were projected into the developmental PC basis, only approximately 40% of cancer variance was explained. This implies that roughly 60% of variation lies outside the developmental subspace. Therefore, malignant cell states cannot be explained solely as *distortions* or *extensions* of developmental trajectories.

**The orthogonal component forms a coherent malignant manifold.** To determine whether the orthogonal component represents *structured* variation, we performed PC analysis on the residual embeddings. The portion of the cancer transcriptome that cannot be explained by development corresponds to the residual $R = \tilde{X}_{\text{can}} - X_{\text{can}}^{\text{recon}}$. By construction, this residual lies in the *orthogonal* complement of the developmental subspace. Each cancer cell is therefore decomposed as

$$\mathbf{x}_{\text{can}} = \mathbf{x}_{\text{can}}^{\text{dev,recon}} + \mathbf{x}_{\text{can}}^{\text{res}},$$

where $\mathbf{x}_{\text{can}}^{\text{dev,recon}}$ represents the reconstruction from the developmental subspace projection and $\mathbf{x}_{\text{can}}^{\text{res}}$ is orthogonal to it. To characterize the structure of this residual, we performed PC analysis on the corresponding centered matrix $R_c = R - \bar{R}$, with $\bar{R}$ being the mean of rows as before (this is what we referred as *residual* PC space later on). The first 10 residual PCs explained approximately 67% of residual variance, and the first 30 explained nearly 88%. Thus, the orthogonal component forms a coherent low-dimensional manifold distinct from the developmental manifold.

**Projection scores.** To quantify the degree to which each cell $\mathbf{x}_i$ aligns with developmental programs, we computed a projection score PSdev defined as

$$\text{PSdev}_i = \frac{\|\mathbf{x}_i^{\text{dev,recon}}\|^2}{\|\mathbf{x}_i^{\text{dev,recon}}\|^2 + \|\mathbf{x}_i^{\text{res}}\|^2}.$$

This quantity measures the fraction of the transcriptional signal of cell $i$ that is explained by the developmental subspace, with values close to 0 or 1 indicating transcriptional states dominated by malignant-specific or developmental programs, respectively. The distribution of the PSdevs (Fig. 1C, main text) illustrates that developmental cells exhibit near-complete alignment with the developmental manifold (median PSdev $\approx 0.98$), whereas malignant cells display substantially lower values (median PSdevs $\approx 0.4$), indicating that most states derive less than half of their embedding from developmental axes.

**Prediction of cancer cell states.** To further assess the information content of each component, we asked how well developmental and residual representations predict cancer cell annotations. We trained a multinomial logistic regression classifier using five-fold cross-validation (`max_iter = 2000`, using the Logistic Regression package within the 'scikit-learn' Python module) and evaluated classification accuracy separately for two annotation levels: cancer cell state (N1–N2 and K1–K6; Fig. 1D, main text) and broader condition labels (ductal, transitional, malignant_ductal) spanning disease progression. For the developmental representation, classifiers were trained on the first 30 coordinates of the projection onto the developmental PC space. For the residual representation, classifiers were trained on the first 30 residual PCs. We also evaluated a combined representation obtained by concatenating the first 30 developmental coordinates and the first 30 residual coordinates.

The developmental component alone predicted cancer cell state with high accuracy ($0.94 \pm 0.01$). Strikingly, the residual component alone achieved comparable performance ($0.94 \pm 0.02$), demonstrating that variation outside the developmental subspace encodes substantial information about malignant identity. For broader condition labels, both components remained informative, although with reduced accuracy (developmental: $0.72 \pm 0.02$; residual: $0.73 \pm 0.02$), consistent with greater heterogeneity at this level of annotation. Combining both representations further improved performance, reaching $0.96 \pm 0.02$ accuracy for cancer state and $0.75 \pm 0.02$ for condition labels. Thus, developmental and residual components capture partially complementary aspects of cancer variation.

These results suggest overall that malignant heterogeneity cannot be fully understood as reactivation of developmental programs. Instead, cancer states appear to partially reuse developmental regulatory axes while simultaneously exploring independent directions of variation absent in normal development. This decomposition allows transcriptional heterogeneity in cancer to be interpreted as the combination of *developmental mimicry* and *malignant innovation*.

#### Developmental lineage-referenced coordinate system

**Definition of developmental lineage centroids and directions.** To define a biologically interpretable coordinate system for epithelial differentiation, we used developmental populations corresponding to foregut progenitors and three terminal epithelial lineages: ductal, acinar, and biliary cells. This selection was not intended to exhaustively represent all developmental populations in the dataset. Rather, it provides a compact reference for comparing malignant epithelial states with major foregut-derived pancreatobiliary epithelial branches (Gittes 2009; Shih et al. 2013).

Each developmental cell $\mathbf{x}_i \in \mathbb{R}^{1\times p}$ (i.e., a row in $\tilde{X}_{\text{dev}}$) was represented by its $n_{\text{PC}}$-dimensional vector,

$$\mathbf{x}_i^{\text{dev}} = \mathbf{x}_i B,$$

where $B \in \mathbb{R}^{p\times n_{\text{PC}}}$ contains the retained PC loading vectors as defined above, with $n_{\text{PC}} = 75$. Developmental cells annotated as foregut, ductal, acinar, or biliary were then selected in this space. For each of these population classes $c$, we computed the centroid

$$\mu_{\text{c}}^{PC} = \frac{1}{n_{\text{c}}} \sum_{i\in \text{c}} \mathbf{x}_i^{\text{dev}}.$$

The four centroid norms were similar: foregut = 5.54 ($n = 3948$), ductal = 5.41 ($n = 4000$), acinar = 5.67 ($n = 3747$), and biliary = 6.29 ($n = 1742$). Thus, the foregut centroid should not be interpreted as the geometric origin of this space. We used foregut as a biologically motivated anchor because it represents an early epithelial state.

We now define lineage-associated directions as *centroid-to-centroid* displacement vectors from foregut to each terminal epithelial state:

$$\mathbf{v}_{\text{ductal}} = \mu_{\text{ductal}}^{PC} - \mu_{\text{foregut}}^{PC},$$

$$\mathbf{v}_{\text{acinar}} = \mu_{\text{acinar}}^{PC} - \mu_{\text{foregut}}^{PC},$$

$$\mathbf{v}_{\text{biliary}} = \mu_{\text{biliary}}^{PC} - \mu_{\text{foregut}}^{PC}.$$

Thus, lineage vectors summarize foregut-to-terminal displacements within the developmental PC-coordinate system. These vectors define a lineage-referenced coordinate system. Because they are *not* constrained to be orthogonal, they do not form a strict orthonormal basis, but instead provide three biologically interpretable developmental directions.

**Branch geometry and robustness.** To assess whether these lineage directions define distinct developmental axes, we computed pairwise angles, $\theta_{ij} = \cos^{-1}(\hat{\mathbf{v}}_i \cdot \hat{\mathbf{v}}_j)$, where $\hat{\mathbf{v}}_i = \mathbf{v}_i/\|\mathbf{v}_i\|$. The observed angles and branch lengths were:

| Pair | Angle (°) | $\|\mathbf{v}_i\|$ | $\|\mathbf{v}_j\|$ |
|---|---|---|---|
| Foregut → Ductal *vs.* Foregut → Acinar | 53.6 | 8.25 | 8.79 |
| Foregut → Ductal *vs.* Foregut → Biliary | 60.5 | 8.25 | 8.65 |
| Foregut → Acinar *vs.* Foregut → Biliary | 56.3 | 8.79 | 8.65 |

Thus, the three lineage directions are clearly separated but not orthogonal. They also have comparable lengths, indicating that the angular structure is not driven by one unusually short or unstable branch. This geometry is consistent with a *fan-like* developmental organization rather than a single linear continuum.

Finally, to evaluate robustness, we repeated the angle calculation after bootstrap resampling cells within each developmental population. The resulting distributions were narrow and centered near the original angle estimates, indicating that the lineage directions are reproducible and not driven by sampling noise (Fig. S6).

**Projection of developmental cells onto lineage directions.** To determine whether the centroid-defined directions correspond to occupied developmental structure rather than endpoint geometry alone, each developmental cell $\mathbf{x}_i^{\text{dev}}$ was decomposed *relative* to each branch direction $\mathbf{v}$:

$$t_i = (\mathbf{x}_i^{\text{dev}} - \mu_{\text{foregut}}^{PC}) \cdot \frac{\mathbf{v}}{\|\mathbf{v}\|}, \qquad d_{\perp,i} = \left\| (\mathbf{x}_i^{\text{dev}} - \mu_{\text{foregut}}^{PC}) - t_i \frac{\mathbf{v}}{\|\mathbf{v}\|} \right\|. \tag{1}$$

We also defined a normalized branch coordinate,

$$t_{\text{frac},i} = \frac{t_i}{\|\mathbf{v}\|},$$

so that $t_{\text{frac},i} \approx 0$ corresponds to the foregut anchor and $t_{\text{frac}} \approx 1$ to the corresponding terminal centroid. The orthogonal distance $d_{\perp,i}$ measures deviation from the straight centroid-to-centroid branch axis.

Developmental cells were broadly distributed along all three lineage directions (Fig. S7A). For each branch, we defined a branch corridor as cells located between foregut and the corresponding terminal state, $0 < t_{\text{frac}} < 1$, and close to the branch axis according to a branch-specific empirical threshold on $d_\perp$. Corridor occupancy was substantial for each lineage: 2305/4000 ductal cells, 2349/3747 acinar cells, and 2641/1742 biliary cells (note that corridors are defined geometrically, so they may include cells from multiple developmental annotations).

We further defined candidate intermediate cells as those within the branch corridor with $0.2 \leq t_{\text{frac}} \leq 0.8$, thereby excluding cells near either endpoint. This identified 610 intermediate cells along the ductal branch, 749 along the acinar branch, and 968 along the biliary branch. These results indicate that the lineage directions are not defined solely by terminal centroids, but are populated by cells spanning root-to-terminal continua.

The branch coordinates were also biologically oriented: foregut cells centered near $t_{\text{frac}} = 0$, whereas terminal populations were near $t_{\text{frac}} = 1$ across branches (Fig. S7A). However, orthogonal distances remained broad (Fig. S7B). For each branch, the expected terminal population was closest to its corresponding axis, but differences between labels were modest, with median $d_\perp$ values typically in the range of approximately 11.5-14.0 embedding units. Thus, developmental organization is directionally structured but not confined to narrow, discrete branch trajectories.

To contextualize these distances, we estimated the typical cell embedding norm to be approximately 13.41, with a coordinate-wise standard deviation of approximately 1.55. Therefore, orthogonal distances of approximately 10–15 are within the natural scale of this representation and do not indicate numerical instability. Instead, they support the interpretation that developmental cells occupy broad lineage-referenced corridors within a continuous, overlapping developmental manifold.

### Cancer-state transition vectors relative to developmental geometry

We also examined whether differences between cancer states follow the developmental directions defined above, and whether those differences are captured by the broader developmental PC subspace $\mathcal{D}$. These are related but distinct questions. The lineage-axis analysis asks whether malignant-state differences point along specific foregut-to-terminal developmental directions. The developmental-subspace analysis asks whether those differences lie within the broader linear space spanned by dominant developmental variation. This section provides the quantitative support for Fig. 2 of the main text.

**Malignant-state centroids and transition vectors in developmental PC coordinates.** For lineage-referenced analyses, malignant-state centroids were computed in the same developmental PC-coordinate system used to define the foregut-to-terminal developmental lineage directions. Each *cancer* cell $\mathbf{x}_{\text{can},i} \in \mathbb{R}^{1\times p}$ (i.e., a row in $\tilde{X}_{\text{can}}$) was represented by its $n_{\text{PC}}$-dimensional vector as before, $\mathbf{x}^{\text{dev}}_{\text{can},i} = \mathbf{x}_{\text{can},i}B$. For each annotated state $s$, we computed a centroid in this developmental PC space:

$$\mu_s^{\text{can,PC}} = \frac{1}{n_s}\sum_{i\in s}\mathbf{x}^{\text{dev}}_{\text{can},i},$$

with $n_s$ the number of cells assigned to state $s$. Centroid-to-centroid displacement vectors between malignant states were then defined as

$$\mathbf{d}^{\text{PC}}_{i\to j} = \mu_j^{\text{can,PC}} - \mu_i^{\text{can,PC}}.$$

We considered explicitly `early` (ductal → transitional), `late` (transitional → malignant_ductal), and `total` (ductal → malignant_ductal) state displacements. These shifts should be interpreted as geometric differences between annotated states, not as directly observed temporal trajectories.

**Alignment with developmental lineage directions.** To test whether cancer state differences follow *canonical* developmental directions, we compared the previous displacement vectors with the three lineage axes defined before. For a malignant displacement vector $\mathbf{d}^{\text{PC}}_{i\to j}$ and a developmental lineage direction $\mathbf{v}_k$, alignment was quantified by cosine similarity:

$$\cos(\mathbf{d}^{\text{PC}}_{i\to j}, \mathbf{v}_k) = \frac{\mathbf{d}^{\text{PC}}_{i\to j}\cdot\mathbf{v}_k}{\|\mathbf{d}^{\text{PC}}_{i\to j}\|\,\|\mathbf{v}_k\|}.$$

Values near 1 indicate strong alignment with a developmental direction, values near 0 indicate little directional relationship, and negative values indicate opposition. Across all displacement vectors, cosine similarities with the three developmental axes were small in magnitude, ranging from -0.249 to 0.0176. Thus, differences between malignant states are not organized as movement along the canonical foregut-to-ductal, foregut-to-acinar, or foregut-to-biliary directions.

**Decomposition into developmental and residual components.** We next asked whether displacement vectors were captured by the broader developmental PC subspace, even if they did not align with the three canonical lineage axes. For each malignant state $s$, we computed a centered CLS-space centroid:

$$\mu_s^{\text{can,CLS}} = \frac{1}{n_s}\sum_{i\in s}\mathbf{x}_{\text{can},i},$$

and the associated centroid-to-centroid displacement vector in centered CLS space,

$$\mathbf{d}^{\text{CLS}}_{i\to j} = \mu_j^{\text{can,CLS}} - \mu_i^{\text{can,CLS}}.$$

We then computed its projection into the developmental subspace as $\mathbf{d}^{\text{dev,recon}}_{i\to j} = \mathbf{d}^{\text{CLS}}_{i\to j}BB^T$, where $B$ is the orthonormal developmental PC loading matrix as before. The residual component outside this developmental subspace is $\mathbf{d}^{\text{res}}_{i\to j} = \mathbf{d}^{\text{CLS}}_{i\to j} - \mathbf{d}^{\text{dev,recon}}_{i\to j}$. We quantified the fraction of squared displacement magnitude captured by the developmental subspace as

$$\mathrm{frac}_{\mathrm{dev}}(\mathbf{d}_{i\to j}) = \frac{\|\mathbf{d}_{i\to j}^{\mathrm{dev,recon}}\|^2}{\|\mathbf{d}_{i\to j}^{\mathrm{CLS}}\|^2},$$

and the residual fraction as

$$\mathrm{frac}_{\mathrm{res}}(\mathbf{d}_{i\to j}) = 1 - \mathrm{frac}_{\mathrm{dev}}(\mathbf{d}_{i\to j}).$$

The developmental fraction ranged from 0.429 to 0.610, indicating that malignant state differences contain measurable components within the broader developmental reference space. However, substantial residual components remained outside this space, with $\mathrm{frac}_{\mathrm{res}}$ ranging from 0.390 to 0.571. Therefore, state differences are not developmentally disconnected, but neither are they fully explained by developmental structure. See the summary of results below and Fig. 2, main text.

| Transition | $\mathrm{frac}_{\mathrm{dev}}$ | $\mathrm{frac}_{\mathrm{res}}$ | foregut→ductal | foregut→acinar | foregut→biliary |
|---|---|---|---|---|---|
| ductal → transitional | 0.429 | 0.571 | 0.008 | -0.249 | 0.035 |
| transitional → malignant_ductal | 0.610 | 0.390 | 0.176 | 0.026 | 0.095 |
| ductal → malignant_ductal | 0.450 | 0.549 | 0.137 | -0.20 | 0.102 |

Importantly, lineage-axis alignment and developmental-subspace decomposition answer different questions. The cosine analysis asks whether a malignant displacement vector, represented in developmental PC-score coordinates, follows one of the canonical foregut-to-terminal lineage directions. The developmental-subspace analysis asks whether a malignant displacement has any component within the broader space of developmental variation, using the original centered CLS embedding space and the projection $BB^T$. Thus, malignant displacement vectors can have measurable developmental-subspace components while still showing weak alignment with ductal, acinar, or biliary developmental axes. This distinction explains why cancer states can be positioned within a developmental coordinate system without their differences being organized as movement along canonical developmental trajectories.

#### Single-cell placement of malignant states within developmental branch geometry

Whereas the preceding section analyzed centroid-to-centroid displacement vectors between malignant states, here we study how *individual* cancer cells are positioned relative to the developmental branch geometry.

**Projection of malignant cells onto developmental lineage directions.** We quantified how each malignant cell, represented in $\mathcal{D}$ space as $\mathbf{x}_{\mathrm{can},i}^{\mathrm{dev}}$, lies along each foregut-to-terminal developmental direction and how far it sits from that branch axis. For each developmental branch $b$, we computed the same quantities defined above: a longitudinal coordinate $t_b$, a normalized longitudinal coordinate $t_{\mathrm{frac},b}$, and an orthogonal distance $d_{\perp,b}$.

Across the ductal, acinar, and biliary reference branches, essentially all cells fell within the foregut-to-terminal interval: 99.99% for the ductal branch and 100% for both the acinar and biliary branches. By contrast, developmental cells showed broader spread beyond the endpoint interval, with 68.6%, 69.9%, and 78.6% of cells falling within $0 \leq t_{\mathrm{frac}} \leq 1$ for the ductal, acinar, and biliary branches, respectively. Thus, malignant cells can be positioned within the developmental coordinate system, but their longitudinal positions are compressed within the interior of the developmental range.

Median malignant positions were intermediate along all three branches: $t_{\text{frac}} = 0.47$, 0.46, and 0.41 for the ductal, acinar, and biliary branches, respectively. Developmental cells showed broader corresponding distributions, with medians of 0.61, 0.55, and 0.46. Malignant cells also had lower median orthogonal distances to the branch axes than developmental cells in the same coordinate system: 10.28 versus 13.06 for the ductal branch, 10.09 versus 13.02 for the acinar branch, and 10.47 versus 13.51 for the biliary branch. Together, these results indicate that malignant cells occupy an interior region of the developmental coordinate scaffold rather than extending beyond developmental endpoints.

**Closest developmental branch assignment.** To summarize branch preference at the single-cell level, each malignant cell was assigned to the developmental branch for which it had the smallest orthogonal distance:

$$\text{branch}_i = \arg \min_{b \in \{\text{ductal,acinar,biliary}\}} d_{\perp,b}(\mathbf{x}^{\text{dev}}_{\text{can},i}).$$

Across all malignant cells, the closest branch was acinar for 41.3%, ductal for 36.6%, and biliary for 22.1%. However, nearest-branch assignment was not equivalent to malignant-state identity. Ductal malignant cells were predominantly closest to the acinar branch (89.7%), malignant ductal cells were most often closest to the ductal branch (63.0%), and transitional cells were distributed across all three branches (37.1% acinar, 34.5% ductal, and 28.3% biliary).

These results show that cancer states have interpretable but non-exclusive positional biases within the developmental coordinate system. In other words, developmental branch geometry helps describe where cells lie, but it does not impose discrete lineage identities on malignant states.

### Relative alignment of malignant state centroids with developmental axes

The single-cell branch assignment above describes how individual malignant cells are distributed relative to developmental axes. Here, we summarize first the position of each malignant state at the centroid level to then evaluate its relation to developmental axes. This analysis supports Fig. 3A of the main text.

For each state $s$, we used the malignant-state centroid in developmental PC space, as defined before, $\mu_s^{\text{can,PC}}$. We then defined a vector from the developmental foregut centroid to the malignant-state centroid:

$$\mathbf{r}_s = \mu_s^{\text{can,PC}} - \mu_{\text{foregut}}^{\text{dev,PC}}.$$

The cosine similarity between this malignant position vector and each developmental lineage direction was computed as

$$a_{sk} = \frac{\mathbf{r}_s \cdot \mathbf{v}_k}{\|\mathbf{r}_s\| \, \|\mathbf{v}_k\|}.$$

This quantity measures how strongly the position of malignant state $s$, *relative* to the foregut anchor, aligns with developmental lineage direction $k$ in developmental PC coordinates. The three alignment scores for each malignant state were normalized by their row sum for visualization in Fig. 3A of the main text:

$$\tilde{a}_{sk} = \frac{a_{sk}}{\sum_k a_{sk}}.$$

The plotted values therefore represent relative developmental-axis preference within each malignant state, rather than an absolute measure of developmental constraint. Note that this centroid-level analysis is *positional*: it asks where malignant states lie relative to developmental axes. It is distinct from the transition-vector analysis in Fig. 2, main text, which asks whether differences between cancer states follow developmental directions.

#### Canonical lineage axes do not account for the full developmental component of malignant cells

The analyses above use two related but distinct developmental references: the three biologically interpretable lineage directions and the broader $\mathcal{D}$ subspace. We therefore asked whether the developmental-coordinate component of malignant-cell embeddings could be explained simply by the span of the three canonical lineage axes.

To test this, we compared malignant-cell reconstructions in the developmental PC space with its reconstruction from the span of the three lineage vectors. Let

$$A = \begin{bmatrix} \mathbf{v}_{\text{ductal}} & \mathbf{v}_{\text{acinar}} & \mathbf{v}_{\text{biliary}} \end{bmatrix}$$

denote the matrix whose columns are the foregut-to-terminal lineage vectors in developmental PC space. Because these lineage directions are not orthogonal, projection onto their span was computed using the Moore-Penrose pseudoinverse.

For each cancer cell, we first expressed its developmental PC coordinate relative to the foregut anchor: $\mathbf{q}_i = \mathbf{x}^{\text{dev}}_{\text{can},i} - \mu^{\text{dev,PC}}_{\text{foregut}}$. Its projection onto the three-axis lineage span is then $\mathbf{q}_{i,\text{axes}} = \mathbf{q}_i A A^{+}$. With this, we quantified the fraction of foregut-relative developmental-coordinate magnitude captured by the three canonical lineage axes as

$$f_{\text{axes}|\text{PC}}(i) = \frac{\|\mathbf{q}_{i,\text{axes}}\|^2}{\|\mathbf{q}_i\|^2}.$$

This axis-span analysis is distinct from projection onto the full developmental PC subspace. Projection onto the full developmental subspace is computed in the original centered CLS embedding space as $\mathbf{x}^{\text{dev,recon}}_{\text{can},i} = \mathbf{x}_{\text{can},i} B B^T$, with corresponding developmental fraction

$$f_{\text{dev}}(i) = \frac{\|\mathbf{x}^{\text{dev,recon}}_{\text{can},i}\|^2}{\|\mathbf{x}_{\text{can},i}\|^2}.$$

This full developmental PC subspace captures about 40.4% of the total centered CLS magnitude of malignant cells, while the three canonical lineage axes capture about 21.6% of the malignant cells' foregut-relative developmental-coordinate displacement. Axis proximity explains only part of the developmental-coordinate structure of malignant cells.

This control clarifies the interpretation of lineage-axis analyses. The three lineage directions provide useful biological coordinates for summarizing malignant-cell position, but they do not exhaust the developmental structure captured by the full PC subspace. Developmentally related malignant variation is therefore distributed across broader modes of normal epithelial variation, while a substantial fraction of cancer-cell variation remains outside the developmental reference space.

#### The transitional malignant state is not a simple geometric intermediate in developmental PC space

Our last analysis inspects whether the transitional malignant state forms a simple geometric intermediate between ductal and malignant_ductal states within developmental PC space $\mathcal{D}$. Let

$$D = \mu^{\mathrm{can,PC}}_{\mathrm{ductal}}, \qquad T = \mu^{\mathrm{can,PC}}_{\mathrm{transitional}}, \qquad M = \mu^{\mathrm{can,PC}}_{\mathrm{malignant\_ductal}}$$

denote the three malignant-state centroids in the PC-score space defined previously. We defined the endpoint-connecting vector $DM = M - D$, and the ductal-to-transitional vector $DT = T - D$. The scalar position of $T$ along the ductal-to-malignant_ductal endpoint axis was

$$t = \frac{DT \cdot DM}{\|DM\|^2},$$

with a computed value of $t = 0.627$, i.e., the orthogonal projection of $T$ falls between the two endpoint centroids. Moreover, the closest point on the endpoint-connecting line is $P = D + tDM$, and the perpendicular deviation from this line was $d_\perp = \|T - P\|$. We can also compute the *relative* perpendicular deviation,

$$d_\perp^{\mathrm{rel}} = \frac{d_\perp}{\|DM\|},$$

and the angle between the ductal-to-transitional and ductal-to-malignant_ductal directions:

$$\theta = \cos^{-1}\left(\frac{DT \cdot DM}{\|DT\|\,\|DM\|}\right).$$

Thus, after collapsing developmental PC space onto the one-dimensional ductal-to-malignant_ductal axis, the transitional state appears intermediate. However, the perpendicular distance from the endpoint-connecting line was $d_\perp = 4.479$, while the ductal-to-malignant_ductal centroid separation was $DM = 6.992$. Therefore, $d_\perp^{\mathrm{rel}} = 0.641$, indicating that the off-axis deviation of the transitional centroid was approximately 64.1% of the endpoint separation. In addition, the ductal-to-transitional direction formed a 45.6° angle with the ductal-to-malignant_ductal direction. These values indicate that the transitional centroid is not collinear with the ductal and malignant_ductal centroids in developmental PC space.

Therefore, the transitional state appears intermediate only after one-dimensional projection onto the ductal-to-malignant_ductal axis. In the full developmental PC-coordinate space, it occupies an off-axis position rather than lying on the direct endpoint-connecting path. This supports the conclusion that the transitional state should not be interpreted as a simple developmental bridge between ductal and malignant_ductal cancer states.

**Summary: Developmental PC analysis defined two related representations.** The analyses in the main text and in the previous sections distinguish three related questions. First, where do cancer cells lie within the developmental coordinate system? Second, are those positions explained by proximity to the canonical foregut-to-terminal lineage axes? Third, do differences between malignant states follow developmental directions or form a simple developmental trajectory? To avoid mixing coordinate systems, lineage-position analyses were performed in the $n_{\mathrm{PC}}$-dimensional developmental PC-score space, whereas developmental-subspace and residual decompositions were performed in developmental-mean-centered CLS space using the same developmental $B$ loading matrix.

1st The retained PC scores provided a 75-dimensional coordinate system for comparing cells along dominant developmental variation. This representation was used for lineage-reference analyses, including developmental lineage centroids, foregut-to-terminal lineage vectors, malignant-state centroids in developmental coordinates, branch-position metrics, and cosine alignment between malignant transitions and developmental lineage directions.

2nd The retained PCs also defined a linear subspace of the original CLS embedding space. Projection onto this subspace followed by subtraction from the developmental-mean-centered CLS embedding, was used to decompose malignant-cell variation into developmental and residual components.

Thus, lineage-coordinate analyses and residual-decomposition analyses used the same developmental PC analysis model but different representations of it.

#### Biological annotation of developmental reference axes

To assign biological meaning to the CLS-derived developmental reference space, we used gene-level correlations and curated mouse gene programs. These programs represent major epithelial developmental and lineage-associated processes (`Supplementary Note 1`). We score the expression of a program as the average expression of the corresponding gene markers.

We annotated two classes of *axes*. First, for each of the first ten PCs, we correlated the coordinate of each cell along that component with the expression of every gene. This produced ranked gene lists for each axis, including genes positively and negatively associated with each PC. Second, we used the lineage-specific directions defined by $(\mathbf{v}_{\text{ductal}}, \mathbf{v}_{\text{acinar}}, \mathbf{v}_{\text{biliary}})$ to project each cell $i$ onto the corresponding unit vector $(\hat{\mathbf{v}}_i = \mathbf{v}_i/\|\mathbf{v}_i\|)$, producing a scalar lineage coordinate (that we named $t_i$ before). Gene-level correlations were computed between this scalar coordinates and expression of each gene.

To summarize axis-level biology, we correlated each PC axis or lineage projection with the curated *per*-cell program scores using Spearman correlation. This generated an axis-by-program correlation matrix (Fig. S8). This annotation revealed that the developmental reference space captured several *interpretable* biological programs. The strongest lineage-specific result was observed for the foregut-to-acinar direction, which showed high positive correlations with both the acinar differentiation program and the secretory digestive enzyme program. The top genes increasing along this direction included acinar-associated markers and regulators such as Pnliprp1, Cpa2, Rbpjl, and Nr5a2, supporting the interpretation that this direction represents acinar lineage progression. The same axis also showed a moderate positive association with proliferation, indicating that acinar differentiation in this representation may be partially coupled to proliferative developmental state.

Among the generic CLS components, `CLS_devPC3` (developmental PC3) showed the strongest program-level annotation. It was highly positively correlated with the proliferation/cell-cycle program and also showed positive associations with epithelial maturation, foregut early epithelial identity, biliary-like differentiation, and ductal differentiation. Its top positively associated genes included proliferation- and chromatin/RNA-associated genes, including Mki67, consistent with a proliferative epithelial or progenitor-like state. Thus, `CLS_devPC3` appears to capture a major proliferative epithelial developmental axis rather than a purely lineage-specific differentiation program.

A second generic component, `CLS_devPC10`, showed negative associations with the acinar differentiation and secretory digestive enzyme programs. This indicates that the negative loading of `CLS_devPC10` is enriched for acinar/secretory biology, whereas the positive loading was associated with genes such as Col4a1, Col4a2, Fn1, Trp63, and Sulf1, suggesting a contrasting extracellular matrix, basal-like, or non-acinar epithelial state. Additional CLS components showed weaker or mixed biological annotations. For example, `CLS_devPC4` was negatively associated with epithelial maturation, ductal differentiation, biliary-like differentiation, and progenitor/fetal-like programs, while `CLS_devPC7` showed only a weak acinar-associated signal. Several components, including `CLS_devPC2` and `CLS_devPC8`, were not strongly explained by the curated programs and were therefore left unassigned.

The foregut-to-ductal and foregut-to-biliary lineage projections showed more modest and mixed biological associations than the foregut-to-acinar direction. The foregut-to-ductal direction correlated positively with fetal/progenitor-like, biliary-like, and ductal differentiation programs, supporting a weak-to-moderate ductal/biliary epithelial interpretation. However, canonical ductal markers were not dominant among the top-ranked genes, and the axis also showed a modest inverse association with the neuroendocrine-like peptide hormone program. Therefore, this direction was interpreted cautiously as a modest ductal/biliary-associated developmental transition rather than a clean ductal differentiation axis. The foregut-to-biliary direction was positively associated with the biliary-like differentiation program and included biologically relevant genes such as Pkhd1, Bicc1, Onecut1, Hnf1b, and Prox1. However, this direction also correlated similarly with acinar and secretory digestive enzyme programs, suggesting that it captures a broader differentiation-away-from-foregut signal rather than a uniquely biliary-specific fate.

Overall, this analysis supports the conclusion that the developmental reference space contains biologically interpretable structure. The most robust and lineage-specific signal corresponds to acinar differentiation, while ductal and biliary-associated structure is present but weaker and less specific. In addition, the learned space captures a prominent proliferative epithelial/progenitor-like axis. By contrast, the neuroendocrine-like peptide hormone program showed only weak or modest associations, indicating that this hormone activity is not a primary axis of variation in this reference space.

#### Residual malignant axes capture lineage-like and cancer-adaptive programs

Annotation of residual cancer CLS axes revealed that malignant variation remaining after reconstruction from the developmental reference basis had a biological foundation (Fig. S9). The leading residual axis, `ResCLS_PC1` (Residual PC1), was characterized by a strong contrast between residual acinar/secretory differentiation and stress-associated malignant programs. The positive loading was enriched for acinar/secretory genes, including Zg16, Rnase1, Cela1, Pnlip, Ctrl, Pla2g1b, Clps, Cela3b, Cela2a, Ctrc, Aqp12, and Cpa2. In contrast, the negative loading was associated with AP-1/immediate early response, KRAS/MAPK signaling, senescence/senescence-associated secretory phenotype (SASP), inflammatory signaling, apoptosis, and epithelial stress programs. Thus, `ResCLS_PC1` primarily separates residual pancreatic acinar identity from a stress-adaptive malignant state.

`ResCLS_PC2` showed the clearest *cancer-adaptive* annotation. Its positive loading was strongly enriched for EMT/invasion and extracellular matrix (ECM)-remodeling programs and additionally associated with senescence/SASP, stemness/progenitor-like identity, hypoxia, glycolytic metabolic rewiring, and therapy-resistance/stress-tolerance signatures. Representative positive-loading genes included Tpm1, Tpm4, Sdc1, Antxr2, Anxa5, Cdkn2b, Fn1, Itgb1, and Cnn3, consistent with a matrix-remodeling and invasive malignant cell state.

`ResCLS_PC4` captured a distinct stress-response versus metabolic/proliferative contrast. The positive loading was enriched for AP-1/immediate early response genes, including Jun, Klf6, Dusp1, Ccn1, Btg2, Fosb, Jund, Junb, Zfp36, Atf3, and Egr1. The opposing loading was marked by mitochondrial respiratory chain genes, including Atp5f1e, Atp5mf, Cox5a, Uqcr11, Tomm7, Ndufa3, Ndufs6, and multiple cytochrome oxidase components, and showed relative enrichment for oxidative phosphorylation and proliferative/metabolic programs. This axis therefore separates an AP-1-driven stress-response state from a metabolically active mitochondrial/proliferative state.

Additional residual axes captured more moderate or mixed malignant programs. `ResCLS_PC3` was associated with epithelial identity, unfolded protein response, AP-1/immediate early response, and KRAS/MAPK signaling, with positive-loading genes including Epcam, Cldn7, Rab25, Tmprss4, and Atp1b1. `ResCLS_PC6` displayed a potential neuroendocrine-like positive loading marked by Scg5, Rab3b, Resp18, Syt13, Neurod1, and Gnao1, opposing a more acinar/epithelial stress-associated loading. `ResCLS_PC8` contrasted epithelial/acinar features with a negative loading enriched for therapy-resistance/stress-tolerance, TGF$\beta$, KRAS/MAPK, interferon response, unfolded protein response, EMT/invasion, and glycolytic rewiring programs. `ResCLS_PC9` separated a ductal-like epithelial loading marked by Krt19, Ctse, Itgb4, Lamb3, and Emp1 from a MYC/UPR/proliferative stress-associated loading. `ResCLS_PC10` separated an epithelial metabolic-stress loading, marked by Cldn18, Krt20, Aldoa, Glul, Muc1, Lgals4, and Gsta4, from a proliferative cell-cycle loading marked by Stmn1, Mad2l1, Birc5, Cks1b, Pbk, Cdca3, and Cdca8.

Together, these results demonstrate that residual malignant variation is not an unstructured remainder after subtraction of the developmental reference component. Instead, the residual cancer space retains biologically coherent structure composed of both residual pancreatic differentiation-like programs and cancer-adaptive states, including EMT/invasion, AP-1 stress signaling, metabolic remodeling, interferon/UPR activity, MYC-associated proliferative stress, and therapy-resistance-associated programs.

#### Annotation of malignant states by developmental alignment and residual cancer programs

To determine whether malignant cell states were defined primarily by their position in the developmental reference space, or whether they also carried residual cancer-associated programs, we generated an integrated biological annotation for each cancer state. This analysis combined four complementary measurements: developmental-lineage alignment, developmental program activity, cancer program activity, and residual cancer-associated axis enrichment.

Cancer cells were first positioned in the developmental coordinate system. Each cell was projected onto the previously defined lineage vectors, yielding scalar developmental-alignment scores along the foregut-to-ductal, foregut-to-acinar, and foregut-to-biliary directions. For each malignant state, these scores were $z$-scored across cancer cells and averaged within state to summarize relative developmental alignment. These alignment summaries are reported in Supplementary Table S1 and visualized at the centroid level in Fig. fig3A, main text.

In parallel, we scored curated developmental gene programs in each cancer cell (`Supplementary Note 1`). Program scores were $z$-scored across cancer cells and averaged within each malignant state, producing a state-by-program matrix of relative developmental program enrichment (Fig. S10A). To quantify cancer-associated biology, we similarly scored curated malignant and adaptive programs in each cancer cell (`Supplementary Note 2`). These program scores were again $z$-scored across cancer cells and averaged within each malignant state to identify cancer programs enriched in each population (Fig. S10B).

Finally, we quantified enrichment of each state along residual cancer-associated axes. Recall that residual axes were computed by projecting CLS representations onto the developmental reference basis, reconstructing the component explained by this developmental basis, and subtracting the reconstructed component from the original representation. The resulting residual vectors were centered across cancer cells and decomposed by PC analysis. These residual PCs represent malignant variation not captured by the linear developmental reference basis. For state-level annotation, residual PC scores were $z$-scored across cancer cells and averaged within each malignant state (Fig. S10C). This linked each malignant state to residual cancer-associated programs, including acinar/secretory identity, AP-

1/KRAS stress, epithelial/acinar versus EMT/stress-adaptive states, glycolytic/hypoxic/oxidative-stress programs, inflammatory/antigen/p53-associated programs, and cell-cycle-associated states.

**Ductal state.** The state initially labeled *ductal* ($n$=5,626 cells, see also PDAC manifold discussion) showed the strongest positive developmental alignment along the foregut-to-acinar axis rather than aligning with ductal differentiation (Supplementary Table S1; Fig. fig3A, main text). It was enriched for acinar differentiation and secretory digestive enzyme programs, while showing low or negative enrichment for neuroendocrine-like peptide hormone and proliferation/cell-cycle programs (Supplementary Fig. S10A).

Its marker genes included acinar and digestive enzyme-associated genes such as Pnlip, Ctrc, Pla2g1b, Zg16, Hsd17b13, Cela1, Try4, Ctrl, Aqp12, and Klk1b3. Consistent with this interpretation, the state showed strong positive enrichment along `ResCLS_PC1`, corresponding to the acinar/secretory versus AP-1/KRAS stress residual axis, and positive enrichment along `ResCLS_PC8`, corresponding to an epithelial/acinar versus EMT/stress-adaptive residual axis (Supplementary Fig. S10C). In contrast, this state showed low enrichment for inflammatory signaling, interferon response, ECM remodeling/invasion, antigen presentation, and cell-cycle programs (Supplementary Fig. S10B).

These results indicate that the state initially labeled *ductal* is better interpreted as an acinar/secretory-like malignant state. The original label should therefore be treated cautiously in downstream interpretation. Biologically, this population appears relatively differentiated and less enriched for adaptive cancer programs than the malignant ductal and transitional states.

**Malignant ductal state.** The *malignant_ductal* state ($n$=8,855 cells) showed the clearest ductal/progenitor-like developmental profile. It was positively aligned with the foregut-to-ductal direction, while showing lower relative alignment with the foregut-to-acinar and foregut-to-biliary developmental directions (Supplementary Table S1; Fig. fig3A, main text). Developmental program analysis showed enrichment for foregut early epithelial identity, morphogenesis/epithelial maturation, ductal differentiation, and proliferation/cell-cycle programs (Supplementary Fig. S10A). Marker genes for this state included 3110039M20Rik, Cdx2, Foxg1, Klra4, Crisp1, Twist1, Nrn1, Krt14, Chst13, Sox17, Ptges, Igf2bp1, Dlk1, and Mgarp, consistent with a ductal/progenitor-like and developmentally plastic malignant epithelial state.

In addition to this developmental alignment, the malignant ductal state showed strong enrichment for cancer-associated metabolic and stress-adaptive programs. These included glycolytic metabolic rewiring, oxidative stress, hypoxia, oxidative phosphorylation, stemness/progenitor-like identity, and MYC targets (Supplementary Fig. S10B). At the residual-axis level, this state was enriched along `ResCLS_PC7`, corresponding to glycolytic/hypoxic/oxidative-stress biology, and `ResCLS_PC5`, corresponding to weak inflammatory/antigen/p53-associated variation. It also showed negative enrichment along `ResCLS_PC1`, indicating movement away from the acinar/secretory residual loading and toward the opposite AP-1/KRAS/stress-associated loading (Supplementary Fig. S10C). Together, these results indicate that the malignant ductal state combines ductal/biliary-like developmental identity with elevated metabolic, proliferative, and stress-adaptive cancer programs.

**Transitional state.** The *transitional* state ($n$=13,650 cells) did not show strong positive enrichment along the foregut-to-ductal, foregut-to-acinar, or foregut-to-biliary axes (Supplementary Table S1; Fig. fig3A, main text). Developmental program scores were close to neutral, with only modest enrichment for neuroendocrine-like peptide hormone, morphogenesis/epithelial maturation, acinar differentiation, and secretory digestive enzyme programs (Supplementary Fig. S10A). This suggests that the transitional population is not well explained by normal developmental lineage programs alone.

The marker genes of the transitional state suggested a distinct biology. Top markers included Gip, Cck, Gast, Fev, Chgb, Sez6, Hepacam2, Rgs13, Prss3, Neurod1, Tph1, and Penk, consistent with neuroendocrine-like or peptide hormone-associated features. Cancer program analysis showed modest enrichment for p53 pathway, AP-1 immediate early response, KRAS/MAPK signaling, ECM remodeling/invasion, unfolded protein response, and antigen-presentation programs (Supplementary Fig. S10B). At the residual-axis level, the transitional state showed negative enrichment along `ResCLS_PC7`, `ResCLS_PC5`, `ResCLS_PC1`, and `ResCLS_PC10`, indicating that its malignant identity is not dominated by the same glycolytic/hypoxic/oxidative-stress or acinar/secretory residual axes that characterize the other states (Supplementary Fig. S10C).

These findings argue against interpreting the transitional state as a faithful normal developmental intermediate. Instead, the transitional state appears to represent a malignant adaptive or transitional cell state with neuroendocrine-like features and modest stress-associated cancer programs. This distinction is important because the transitional population occupies neither a strongly ductal nor strongly acinar developmental program state, yet it carries malignant features that distinguish it from a simple developmental intermediate.

Together, these analyses show that malignant state identity is shaped by both developmental-like positioning and residual cancer-associated programs. The state initially labeled *ductal* was dominated by acinar/secretory features and showed relatively low activation of adaptive cancer programs. The *malignant ductal* state combined ductal/biliary and progenitor-like developmental alignment with strong metabolic and stress-associated malignant programs. The *transitional* state showed limited evidence of normal developmental program enrichment and instead displayed neuroendocrine-like markers together with modest AP-1/KRAS, p53, ECM remodeling, unfolded protein response, and antigen-presentation activity. Thus, cancer state identity is not explained solely by developmental coordinate position. In particular, the transitional state is better interpreted as a cancer-associated adaptive state rather than a normal developmental intermediate.

#### Malignant state-displacement vectors are associated with cancer-adaptive reprogramming

We next annotated the biological programs associated with malignant state-displacement vectors. For each pair of malignant states, we defined a centroid-to-centroid displacement vector and projected each cancer cell onto that vector. These displacement vectors were computed in both *developmental* PC space and *residual* cancer PC space.

The *ductal-to-transitional transition*, which starts from an acinar/secretory-like malignant state, was marked by coherent biological program changes like loss of acinar digestive enzyme genes, including Zg16, Cela1, Rnase1, Pnlip, Pla2g1b, Ctrl, Try4, Clps, Cela3b, Cela2a, Prss3b, Pdia2, Ctrc, Aqp12, and Cpa2. Programs increasing toward the transitional state included ductal differentiation, epithelial morphogenesis/maturation, foregut early epithelial identity, biliary-like differentiation, and proliferation/cell-cycle programs (Fig. S11A). Cancer-associated programs increasing toward the transitional state included AP-1 immediate early response, KRAS/MAPK signaling, p53 pathway, epithelial identity, inflammatory signaling, senescence/SASP, and unfolded protein response (Fig. S11B). Thus, the ductal-to-transitional transition captures loss of differentiated secretory identity and acquisition of epithelial stress-response programs.

The transition from the *transitional to the malignant ductal* state was associated with increas-

| Cancer state | Developmental alignment | Representative marker genes | Enriched programs | Interpretation |
|---|---|---|---|---|
| *ductal* | Strong foregut-to-acinar alignment; enriched acinar differentiation and secretory digestive enzyme programs | Pnlip, Ctrc, Pla2g1b, Zg16, Hsd17b13, Cela1, Try4, Ctrl, Aqp12, Klk1b3 | Acinar differentiation and secretory digestive enzymes; residual acinar/secretory enrichment; low inflammatory, ECM/invasion, interferon, antigen-presentation, and cell-cycle activity | Acinar/secretory-like malignant state; original label should be interpreted cautiously |
| *malignant ductal* | Foregut-to-ductal alignment; enriched foregut epithelial, ductal, morphogenesis/maturation, and proliferative programs | 3110039M20Rik, Cdx2, Foxg1, Klra4, Crisp1, Twist1, Nrn1, Krt14, Chst13, Sox17, Ptges, Igf2bp1, Dlk1 | Glycolytic rewiring, oxidative stress, hypoxia, oxidative phosphorylation, stemness/progenitor-like identity, MYC targets, and residual glycolytic/oxidative-stress enrichment | Ductal/progenitor-like malignant state with metabolic and stress-adaptive programs |
| *transitional* | No strong enrichment for canonical ductal, acinar, or biliary developmental axes | Gip, Cck, Gast, Fev, Chgb, Sez6, Hepacam2, Rgs13, Prss3, Neurod1, Tph1, Penk | Modest p53 pathway, AP-1 immediate early response, KRAS/MAPK signaling, ECM remodeling, unfolded protein response, and antigen-presentation activity | Malignant adaptive/transitional state with neuroendocrine-like features; not a faithful normal developmental intermediate |

Table S1: Biological interpretation of malignant states based on developmental alignment, marker genes, cancer program enrichment, and residual cancer-axis contributions.

ing metabolic and proliferative malignant programs. Genes increasing toward the malignant ductal endpoint included Gapdh, Wfdc2, Aprt, S100a6, Tpi1, Pkm, Tomm20, Capg, Pgam1, and Cdkn2a. Developmental programs associated with this transition included decreased acinar differentiation, decreased secretory digestive enzyme activity, decreased neuroendocrine-like peptide hormone signal, and increased progenitor/fetal-like and proliferation/cell-cycle programs (Fig. S11A). Cancer programs increasing toward the malignant ductal state included glycolytic metabolic rewiring, oxidative stress, hypoxia, EMT/invasion, cell cycle, stemness/progenitor-like identity, and MYC targets (Fig. S11B). These findings indicate that movement from the transitional state to the malignant ductal state involves acquisition of a metabolically rewired, hypoxic, invasive, and proliferative malignant phenotype.

The full ductal-to-malignant_ductal transition captured the dominant malignant reprogramming trajectory. This transition was associated with strong loss of acinar differentiation and secretory digestive enzyme programs, together with gain of foregut early epithelial identity, epithelial morphogenesis/maturation, ductal differentiation, progenitor/fetal-like, and proliferation/cell-cycle programs (Fig. S11A). At the cancer-program level, movement toward the malignant ductal endpoint was associated with stemness/progenitor-like identity, glycolytic rewiring, oxidative stress, epithelial identity, MYC targets, senescence/SASP, hypoxia, EMT/invasion, apoptosis, TGF-beta signaling, therapy-resistance/stress-tolerance, oxidative phosphorylation/mitochondrial programs, and cell-cycle activity (Fig. S11B). These patterns were observed for transition vectors defined in both developmental coordinate space and residual cancer space, indicating that the biological interpretation of the transition is robust to the coordinate system used to define the state-centroid vector.

Together, these results show that malignant state differences are not normal developmental transitions. Instead, they describe cancer-adaptive reprogramming in which differentiated acinar/secretory identity is lost and ductal/progenitor-like epithelial, metabolic, hypoxic, oxidative-stress, MYC, inflammatory, senescence, unfolded-protein-response, proliferative, and invasive programs are gained. The transitional state appears to occupy an intermediate malignant reprogramming position characterized by AP-1/KRAS, p53, inflammatory, senescence/SASP, and unfolded protein response activity, whereas the malignant ductal endpoint is characterized by stronger glycolytic rewiring, hypoxia, oxidative stress, stemness/progenitor-like identity, MYC activity, EMT/invasion, and proliferation.

#### Supplementary Note 1: Mouse transcriptional programs

To assist interpretation of mouse epithelial states, we defined a curated set of marker-based transcriptional programs capturing major developmental, differentiation, and functional axes. These programs were not intended to represent mutually exclusive cell identities. Instead, they provide compact gene-set summaries that can be scored across cells to aid biological annotation. We performed an enrichment-based validation to further assess their biological consistency.

For each program, the component genes were queried against public functional, pathway, tissue, cell-type, and phenotype annotation resources, including Gene Ontology, KEGG, Reactome, Mouse Gene Atlas, PanglaoDB, and mammalian phenotype-related annotations. Enriched terms from the queried libraries were then compared with program-specific expected biological keywords. In addition to testing the full curated programs, we also evaluated *stricter* diagnostic subprograms that separated shared epithelial, ductal, acinar, progenitor/fetal-like, and neuroendocrine components. These diagnostic subprograms were used to determine whether the full programs were supported by the intended biological component or by a more generic marker backbone. All full programs showed significant annotation support for their assigned biological labels; however, gene-set overlap and term-level inspection identified several cases where cautious interpretation was required.

The *foregut early epithelial identity* program captures an early epithelial state with foregut/endoderm-associated transcriptional features. It combines transcription factors such as Sox2, Sox9, Foxa1, Foxa2, and Hnf1b with epithelial structural and junctional genes including Krt8, Krt18, Epcam, and Cldn4. The full program was strongly supported by epithelial cell-type annotations, whereas the stricter transcription-factor subprograms showed support for primitive endoderm-related annotations, including terms driven by Hnf1b and Foxa2. Thus, high scores for the full program should be interpreted as evidence of an epithelial state with potential early foregut/endoderm-like features, particularly when the transcription factor component is co-expressed with the epithelial marker component. The program should not be interpreted as definitive foregut identity based on epithelial markers alone.

The *ductal differentiation* and *biliary-like differentiation* programs are related but not independent. Both contain a shared ductal epithelial backbone, including markers such as Krt7, Krt19, Sox9, Muc1, Epcam, and Hnf1b. The ductal differentiation program was strongly supported by ductal cell annotations and includes genes associated with ductal epithelial identity and transport, such as Cftr, Slc4a4, Cldn10, and Mmp7. The biliary-like program also showed strong ductal enrichment, but its stricter biliary-like core was supported by cholangiocyte-associated annotations involving genes such as Tff1, Tff2, Slc4a2, and Cldn4. Because the ductal and biliary-like programs share a substantial marker backbone, elevated biliary-like scores are interpreted as a ductal/biliary-like or mucinous ductal epithelial phenotype, rather than as evidence of bona fide biliary lineage conversion.

The *acinar differentiation* and *secretory digestive enzyme* programs both capture exocrine acinar biology and are highly overlapping. They are dominated by digestive enzyme genes, including trypsins, chymotrypsins, elastases, carboxypeptidases, lipase/co-lipase, and amylases. Enrichment analysis strongly supported acinar cell, pancreatic secretion, protein digestion, and digestive enzyme-related annotations. Because the two gene sets showed high overlap, they are best interpreted as closely related acinar/exocrine

| Program | Biological Interpretation | Representative genes |
|---|---|---|
| `foregut_early_epithelial_identity` | Early epithelial/endodermal or foregut-like identity, combining developmental transcription factors with epithelial structural and junctional markers. | Sox2, Sox9, Foxa1, Foxa2, Hnf1b, Krt8, Krt18, Epcam, Cldn4 |
| `ductal_differentiation` | Pancreatic ductal epithelial differentiation, including keratins, ductal transcriptional regulators, mucin-associated genes, and ion transport features. | Krt19, Krt7, Sox9, Cftr, Muc1, Mmp7, Slc4a4, Anxa4, Cldn10, Hnf1b |
| `acinar_differentiation` | Exocrine acinar differentiation and maturation, dominated by digestive enzyme and secretory-effector genes. | Prss1, Prss2, Cela2a, Cela3b, Cpa1, Cpa2, Cpb1, Ctrb1, Ctrb2, Reg1, Reg2, Pnlip, Clps, Amy2a5, Amy2a2 |
| `biliary_like_differentiation` | Ductal/biliary-like epithelial state with mucinous and trefoil-factor features; should be interpreted as a biliary-like transcriptional phenotype rather than definitive lineage conversion. | Krt7, Krt19, Sox9, Epcam, Muc1, Tff1, Tff2, Slc4a2, Anxa4, Cldn4, Cldn10, Hnf1b |
| `proliferation_cell_cycle` | Cycling and mitotic state. This program is orthogonal to lineage identity and should be controlled for when interpreting differentiation trajectories. | Mki67, Top2a, Pcna, Mcm2, Mcm3, Mcm4, Mcm5, Mcm6, Mcm7, Cdk1, Ccnb1, Ccnb2, Ube2c, Birc5 |
| `morphogenesis_epithelial_maturation` | Epithelial organization, adhesion, polarity, tight-junction formation, and basement-membrane-associated maturation. | Epcam, Krt8, Krt18, Krt19, Cdh1, Cldn3, Cldn4, Ocln, Tjp1, Itga6, Lamc2 |
| `secretory_digestive_enzyme_program` | Functional secretory enzyme-production program characteristic of mature exocrine/acinar states. | Prss1, Prss2, Cela2a, Cela3b, Cpa1, Cpa2, Cpb1, Ctrb1, Ctrb2, Pnlip, Clps, Amy2a5, Amy2a2 |
| `progenitor_fetal_like_program` | Developmentally plastic, fetal-like, or progenitor-enriched state. Because it contains epithelial, mucinous, and mesenchymal-associated genes, it should be interpreted alongside other lineage programs. | Sox9, Prom1, Aldh1a1, Aldh1a3, Nes, Muc6, Tff1, Tff2, S100a10, Lgals1, Vim |
| `neuroendocrine_like_peptide_hormone` | Neuroendocrine-like or enteroendocrine-like state, combining secretory-granule machinery, endocrine transcription factors, and peptide hormone genes. | Chga, Chgb, Syp, Scg2, Scg3, Scg5, Snap25, Syt1, Syt4, Syt13, Rab3a, Rab3b, Neurod1, Fev, Isl1, Pax6, Nkx2-2, Gip, Cck, Gast, Tph1, Penk, Resp18 |

Table S2: Curated mouse transcriptional programs used for epithelial-state annotation.

enzyme programs rather than as fully distinct biological states. The acinar differentiation program is slightly broader because it includes Reg1 and Reg2, whereas the secretory digestive enzyme program more directly captures the functional enzyme-producing component of the acinar state.

The *proliferation/cell-cycle* program is included as an orthogonal state variable rather than as a lineage program. It contains canonical cycling and mitotic genes such as Mki67, Top2a, Pcna, Mcm2, Mcm3, Mcm4, Mcm5, Mcm6, Mcm7, Cdk1, Ccnb1, Ccnb2, Ube2c, and Birc5. The enrichment-based validation strongly supported cell-cycle, DNA-replication, and mitotic annotations for this program. Accordingly, this score should be interpreted separately from differentiation identity, because cycling cells may transiently show altered lineage-program scores depending on cell-cycle state.

The *morphogenesis epithelial maturation* program captures epithelial organization, adhesion, junctional maturation, and tissue architecture. It includes epithelial structural genes, adherens-junction and tight-junction components, integrin-associated genes, and basement membrane-associated genes, including Epcam, Krt8, Krt18, Krt19, Cdh1, Cldn3, Cldn4, Ocln, Tjp1, Itga6, and Lamc2. The full program was supported by epithelial cell-type annotations, and the stricter junctional subprogram was supported by cell-cell junction terms. This score is therefore useful for identifying epithelial maturation, junctional organization, and tissue architecture, but it should be distinguished from lineage-specific differentiation programs such as acinar, ductal, or neuroendocrine-like programs.

The *progenitor/fetal-like* program marks a less mature, developmentally plastic, or fetal-like epithelial state, but it should not be treated as a pure progenitor signature. It combines progenitor-associated genes such as Sox9, Prom1, Aldh1a1, Aldh1a3, and Nes with fetal-like, mucinous, and plasticity-associated markers including Tff1, Tff2, S100a10, Lgals1, and Vim. The enrichment-based validation supported epithelial and developmental annotations, but the strongest expected term for the full program involved epithelial structure maintenance and was driven by Tff1 and Tff2. In addition, the diagnostic subprograms indicated that the full program contains fetal/ mucinous and mesenchymal/plasticity-associated components. Elevated scores should therefore be interpreted cautiously as a progenitor/fetal-like or plasticity-associated epithelial state, ideally in combination with epithelial, ductal, acinar, proliferation, and mesenchymal/plasticity-associated scores.

Finally, the *neuroendocrine-like peptide hormone* program captures a neuroendocrine or enteroendocrine-like transcriptional axis. It includes secretory granule and synaptic vesicle genes such as Chga, Chgb, Scg2, Scg3, Scg5, Snap25, Syt1, Syt4, Syt13, Rab3a, Rab3b, and Resp18, together with endocrine transcription factors and peptide hormone genes including Neurod1, Fev, Isl1, Pax6, Nkx2-2, Gip, Cck, Gast, Tph1, and Penk. The enrichment-based validation strongly supported enteroendocrine/neuroendocrine-related annotations, with the top cell-type enrichment involving enteroendocrine cells and multiple hormone, transcription factor, and secretory machinery genes. This program is therefore best interpreted as evidence for a neuroendocrine-like or enteroendocrine-like state, particularly when secretory machinery, endocrine transcription factors, and peptide hormone genes are coherently enriched.

#### Supplementary Note 2: Mouse cancer-associated transcriptional programs

We curated a panel of mouse cancer-associated transcriptional programs to annotate recurrent malignant-cell states and stress-response axes. These programs are again intended as modular gene sets for scoring single cells rather than as mutually exclusive cell-type labels. We similarly evaluated whether the curated labels were consistent with established biological annotations with enrichment-based validation using public annotation resources (described before). Enriched terms were compared with program-specific expected biological keywords, and gene-set overlap was used to identify programs that should be interpreted cautiously. This analysis supported the biological annotation of the curated cancer programs, including strong enrichment for expected terms such as EMT, luminal epithelial identity, TNF/NF-$\kappa$B signaling, interferon signaling, hypoxia, reactive oxygen species response, unfolded protein response, RAS–MAPK signaling, MYC targets, p53 signaling, cell cycle, DNA repair, glycolysis, oxidative phosphorylation, extracellular matrix organization, antigen presentation, TGF-$\beta$ signaling, apoptosis, senescence-associated inflammatory programs, and AP-1/immediate-early response.

Several programs intentionally overlap because core cancer-associated genes participate in multiple biological processes. For example, the `emt_invasion` and `ecm_remodeling_invasion` programs showed substantial overlap, consistent with shared matrix-interaction, collagen, integrin, and protease-associated biology. Similarly, `hypoxia` and `glycolysis_metabolic_rewiring` share metabolic genes such as Slc2a1, Ldha, Pdk1, Eno1, and Aldoa, reflecting the coupling between low-oxygen stress and glycolytic adaptation. The `KRAS_mapk_signaling` and `ap1_immediate_early_response` programs should also be interpreted as related, because AP-1 and immediate-early transcriptional responses can represent downstream MAPK output or acute stimulation states.

The collection covers major tumor-intrinsic phenotypes including epithelial identity, EMT/invasion, KRAS–MAPK signaling, MYC activity, p53 pathway activation, proliferation, DNA damage repair, metabolic rewiring, oxidative phosphorylation, apoptosis, senescence, stemness/progenitor-like features, and therapy-resistance-associated stress tolerance. It also includes programs reflecting tumor–immune interaction and inflammatory state, including NF-$\kappa$B/inflammatory signaling, interferon response, MHC-I antigen presentation, and MHC-II antigen presentation. These immune-associated programs should be interpreted carefully in single-cell data, because inflammatory and MHC-II signals may reflect tumor-intrinsic transcriptional states, immune or stromal contamination, doublets, or microenvironmental exposure. In particular, MHC-II-associated scores should not be considered tumor-intrinsic without independent evidence of malignant-cell identity.

Stress-adaptation modules such as hypoxia, oxidative stress, unfolded protein response, AP-1 immediate-early response, and TGF-$\beta$ signaling capture common environmental and treatment-associated pressures experienced by malignant cells. The `therapy_resist_stress_tolerance` program was supported by annotations related to ABC transporters, stress response, anti-apoptotic buffering, and detoxification; however, this score should be interpreted as therapy-resistance-associated stress tolerance rather than as direct evidence of measured drug resistance in the absence of treatment-response or functional validation.

Table S3: Mouse cancer-associated transcriptional programs used for malignant-state annotation.

| **Program** | **Biological interpretation** | **Representative genes** |
|---|---|---|
| `emt_invasion` | Mesenchymal transition, motility, invasion, matrix interaction, and stromal-like remodeling. | Vim, Fn1, Zeb1, Zeb2, Snai1, Snai2, Twist1, Cdh2, Itga5, Itgav, Itgb1, Mmp2, Mmp9, Mmp14, Col1a1, Col1a2, Col3a1, Sparc, Tnc, Lox, Postn |
| `epithelial_identity` | Epithelial differentiation, epithelial adhesion, cytokeratin expression, and junctional integrity. | Epcam, Cdh1, Krt8, Krt18, Krt19, Krt7, Cldn3, Cldn4, Cldn7, Ocln, Tjp1, Muc1 |
| `inflammatory_signaling` | NF-$\kappa$B-associated inflammatory signaling, cytokine production, chemokine signaling, and inflammatory adhesion. | Nfkb1, Nfkb2, Rela, Relb, Tnf, Tnfaip3, Il1a, Il1b, Il6, Cxcl1, Cxcl2, Cxcl10, Ccl2, Ccl5, Ptgs2, Socs3, Icam1 |

Continued on next page

| Program | Biological interpretation | Representative genes |
|---|---|---|
| interferon_response | Type I/II interferon response, antiviral-like signaling, STAT/IRF activation, and interferon-stimulated genes. | Stat1, Stat2, Irf1, Irf7, Irf9, Isg15, Ifit1, Ifit2, Ifit3, Ifitm1, Ifitm2, Ifitm3, Oas1a, Oas1g, Oas2, Oas3, Mx1, Rsad2, Usp18, Dhx58 |
| hypoxia | Hypoxia adaptation, HIF-associated transcription, angiogenic signaling, anaerobic metabolism, and low-oxygen stress. | Hif1a, Vegfa, Slc2a1, Ldha, Pdk1, Eno1, Aldoa, Bnip3, Ndrg1, Adm, Car9, Serpine1, Ero1a, Ankrd37 |
| oxidative_stress | NRF2-linked antioxidant defense, glutathione metabolism, detoxification, thioredoxin activity, and iron handling. | Nfe2l2, Keap1, Hmox1, Nqo1, Gclc, Gclm, Gsr, Gpx2, Gpx4, Prdx1, Txn1, Txnrd1, Srxn1, Sqstm1, Ftl1, Fth1 |
| unfolded_protein_response | Endoplasmic-reticulum stress, chaperone activity, PERK/ATF/XBP1 signaling, and protein-folding pressure. | Hspa5, Hsp90b1, Ddit3, Atf3, Atf4, Atf6, Xbp1, Ern1, Eif2ak3, Pdia3, Pdia4, Pdia6, Dnajb9, Herpud1, Derl3, Manf, Creld2 |
| KRAS_mapk_signaling | RAS–MAPK output, ERK-responsive transcription, feedback phosphatases, AP-1 activity, and growth-factor signaling. | Dusp4, Dusp5, Dusp6, Etv1, Etv4, Etv5, Spry1, Spry2, Spry4, Fos, Fosl1, Jun, Junb, Egr1, Egr2, Ets1, Myc, Mapk1, Mapk3 |
| myc_targets | MYC-associated biosynthesis, ribosome/nucleolar activity, nucleotide metabolism, protein translation, and growth. | Myc, Max, Mxd1, Npm1, Ncl, Odc1, Cad, Nme1, Nme2, Hspd1, Hspe1, Tfrc, Slc2a1, Phb, Pa2g4, Hnrnpa1, Hnrnpa2b1, Eif4a1 |
| p53_pathway | p53 signaling, cell-cycle arrest, apoptosis priming, DNA-damage response, and stress-induced transcription. | Trp53, Cdkn1a, Mdm2, Bax, Bbc3, Pmaip1, Gadd45a, Gadd45g, Sesn1, Sesn2, Ddb2, Phlda3, Zmat3, Rrm2b, Ccng1 |
| cell_cycle | Proliferation, DNA replication, mitosis, chromosome segregation, spindle assembly, and cycling malignant states. | Mki67, Top2a, Pcna, Mcm2, Mcm3, Mcm4, Mcm5, Mcm6, Mcm7, Cdk1, Ccnb1, Ccnb2, Cenpa, Cenpe, Cenpf, Tpx2, Bub1b, Ube2c, Aspm, Nusap1, Kif11, Kif20b |
| dna_damage_repair | Homologous recombination, mismatch repair, checkpoint signaling, Fanconi anemia pathway, and double-strand break repair. | Brca1, Brca2, Rad51, Rad51ap1, Msh2, Msh6, Pms2, Pcna, Rpa1, Rpa2, Chek1, Chek2, Atm, Atr, Fanca, Fancc, Xrcc5, Xrcc6 |
| glycolysis_metabolic_rewiring | Glucose uptake, glycolytic flux, lactate production, pyruvate handling, and Warburg-like metabolic rewiring. | Slc2a1, Hk1, Hk2, Pfkl, Pfkp, Aldoa, Gapdh, Pgam1, Eno1, Pkm, Ldha, Pdk1, Slc16a3 |

Continued on next page

| Program | Biological interpretation | Representative genes |
|---|---|---|
| oxidative_phospho_mitochondrial | Mitochondrial respiratory-chain activity, TCA-linked respiration, electron transport, and ATP synthase expression. | Ndufa2, Ndufa4, Ndufb3, Ndufb5, Sdha, Sdhb, Uqcrc1, Uqcrc2, Cyc1, Cox4i1, Cox5a, Atp5f1a, Atp5f1b, Atp5mc1 |
| ecm_remodeling_invasion | Matrix deposition, collagen remodeling, integrin signaling, protease activity, and invasive extracellular-matrix interaction. | Fn1, Col1a1, Col1a2, Col3a1, Col4a1, Col4a2, Col5a1, Col5a2, Sparc, Tnc, Postn, Lox, Loxl2, Mmp2, Mmp9, Mmp14, Itga5, Itgav, Itgb1, Vcan, Thbs1 |
| antigen_presentation_mhc_i | MHC-I antigen processing and presentation, immunoproteasome activity, and cytotoxic T-cell visibility. | B2m, H2-K1, H2-D1, Tap1, Tap2, Tapbp, Psmb8, Psmb9, Psmb10, Nlrc5 |
| antigen_presentation_mhc_ii | MHC-II antigen presentation, class-II transcriptional regulation, and antigen-loading machinery. | H2-Aa, H2-Ab1, H2-Eb1, Cd74, Ciita, H2-DMa, H2-DMb1, H2-Ob |
| stemness_progenitor_like | Progenitor-like differentiation state, tumor-initiating features, epithelial plasticity, and partial mesenchymal/stem-like identity. | Sox9, Prom1, Aldh1a1, Aldh1a3, Lgr5, Krt19, Epcam, Tacstd2, Muc1, Nes, Vim, Lgals1, S100a10 |
| therapy_resist_stress_tolerance | Drug efflux, anti-apoptotic signaling, aldehyde metabolism, heat-shock response, antioxidant defense, and stress tolerance. | Aldh1a1, Aldh1a3, Abcb1a, Abcb1b, Abcc1, Abcc3, Abcg2, Bcl2, Bcl2l1, Mcl1, Hspa1a, Hspa1b, Hspa5, Hmox1, Nqo1, Sod2, Gpx2, Lgals1, Tgfb1, Vim |
| tgfb_signaling | TGF-$\beta$ pathway activity, SMAD signaling, EMT induction, matrix-associated signaling, and cytostatic/plasticity responses. | Tgfb1, Tgfb2, Tgfbr1, Tgfbr2, Smad2, Smad3, Smad4, Serpine1, Id1, Id2, Id3, Skil, Snai1, Snai2 |
| apoptosis | Intrinsic and extrinsic apoptotic machinery, caspase activation, mitochondrial apoptosis, and anti-apoptotic buffering. | Bax, Bak1, Bbc3, Pmaip1, Casp3, Casp7, Casp8, Casp9, Apaf1, Bid, Bad, Bcl2, Bcl2l1, Mcl1 |
| senescence_sasp | Cell-cycle arrest, senescence-associated inflammatory secretion, matrix remodeling, and stress-induced paracrine signaling. | Cdkn1a, Cdkn2a, Trp53, Il6, Cxcl1, Cxcl2, Ccl2, Serpine1, Mmp3, Mmp9, Igfbp3, Igfbp5, Gadd45a |
| ap1_immediate_early_response | Immediate-early transcriptional response, AP-1 activation, MAPK feedback, stress response, and acute stimulation signatures. | Fos, Fosb, Fosl1, Fosl2, Jun, Junb, Jund, Atf3, Egr1, Egr2, Dusp1, Dusp5, Dusp6, Ier2, Csrnp1, Zfp36, Btg2, Ccn1, Klf6 |

#### Supplementary Figures

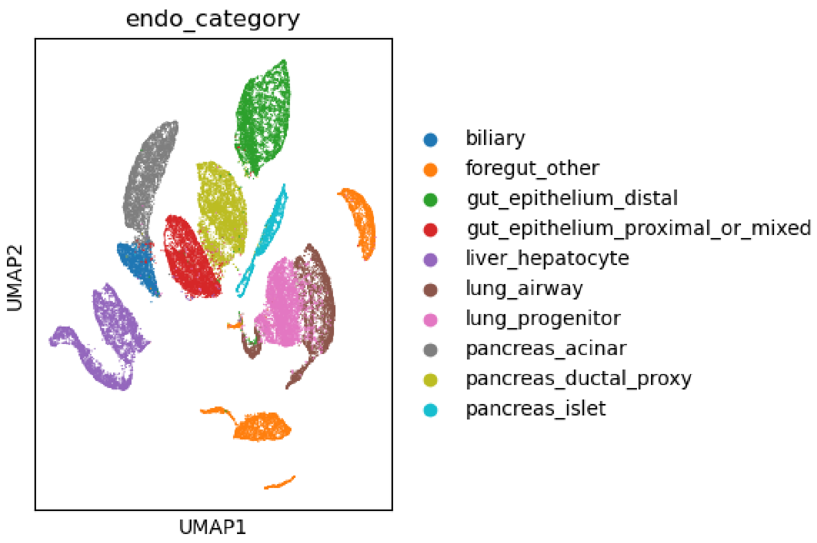


Figure S1: **UMAP representation of the curated epithelial reference atlas derived from the mouse developmental dataset of Qiu** *et al.* Cells belonging to endoderm-derived epithelial tissues were extracted from the E8–P0 developmental atlas and embedded using PCA followed by UMAP. Each point represents a single cell colored by *coarse* epithelial category. Distinct clusters correspond to pancreatic acinar, ductal-proxy, and endocrine cells; hepatobiliary epithelium (hepatocytes and biliary cells); proximal and distal intestinal epithelium; lung progenitor and airway/alveolar epithelium; and foregut-associated epithelial organs (thyroid, parathyroid, thymus). The clear separation of organ-specific populations and the continuity within related epithelial lineages demonstrate that the curated atlas preserves biologically meaningful relationships across foregut and gut epithelia.

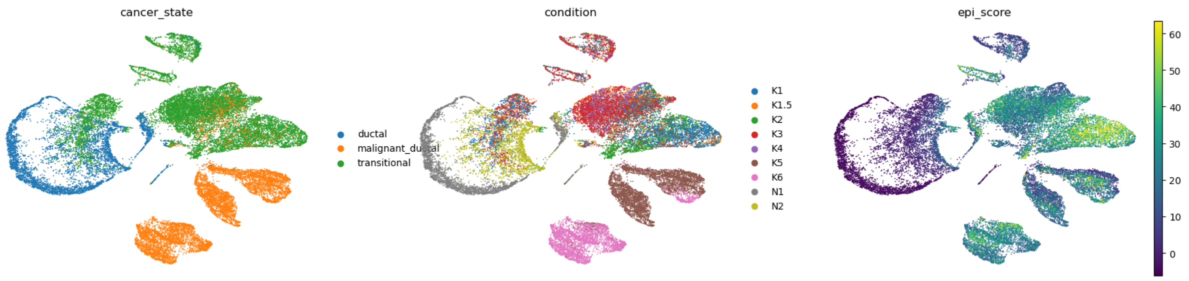


Figure S2: **Transcriptional landscape of epithelial cells across pancreatic tumor progression.** UMAP embedding of the full epithelial dataset from the PDAC progression cohort (Burdziak et al. 2023), including both lineage-traced epithelial cells (mKate2$^+$) and epithelial-enriched samples (EPI). Each point represents a single cell. Left: cells colored by inferred transcriptional phenotype (ductal, transitional, or malignant_ductal). Center: cells colored by experimental condition corresponding to successive stages of pancreatic tumorigenesis, including normal pancreas (N1), injury-induced regeneration (N2), early KRAS-driven metaplasia and preneoplasia (K1–K2), premalignant pancreatic intraepithelial neoplasia (K3–K4), invasive pancreatic ductal adenocarcinoma (K5), and metastatic disease (K6). Right: epithelial gene signature score projected onto the same embedding. The manifold reveals a continuous trajectory linking normal ductal epithelium to preneoplastic and malignant states, with invasive and metastatic tumor cells forming transcriptionally distinct domains enriched for malignant ductal programs.

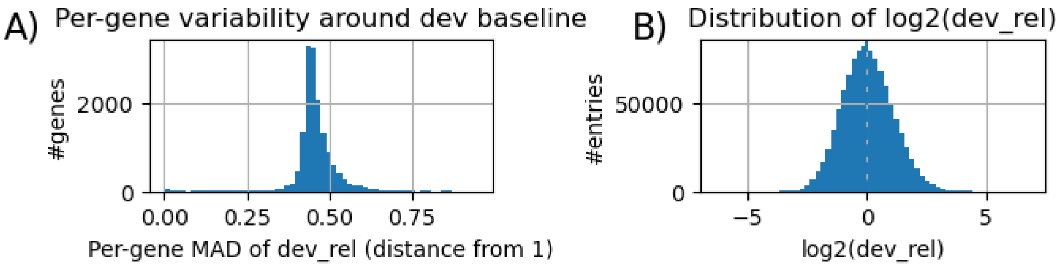


Figure S3: **Valdidation of relative normalizaion.** A) Distribution of *per*-gene median absolute deviations (MADs) of `dev_rel`, quantifying how strongly each gene's expression typically deviates from the developmental baseline (value 1). The 'typical gene' deviates about ±50% from developmental baseline in detected cells. Higher MAD values indicate greater variability across detected cells. B) Histogram of all positive `dev_rel` values shown on a $\log_2$ scale, illustrating the global distribution of relative expression values across genes and cells. The dashed line marks $\log_2(1) = 0$, corresponding to no deviation from baseline.

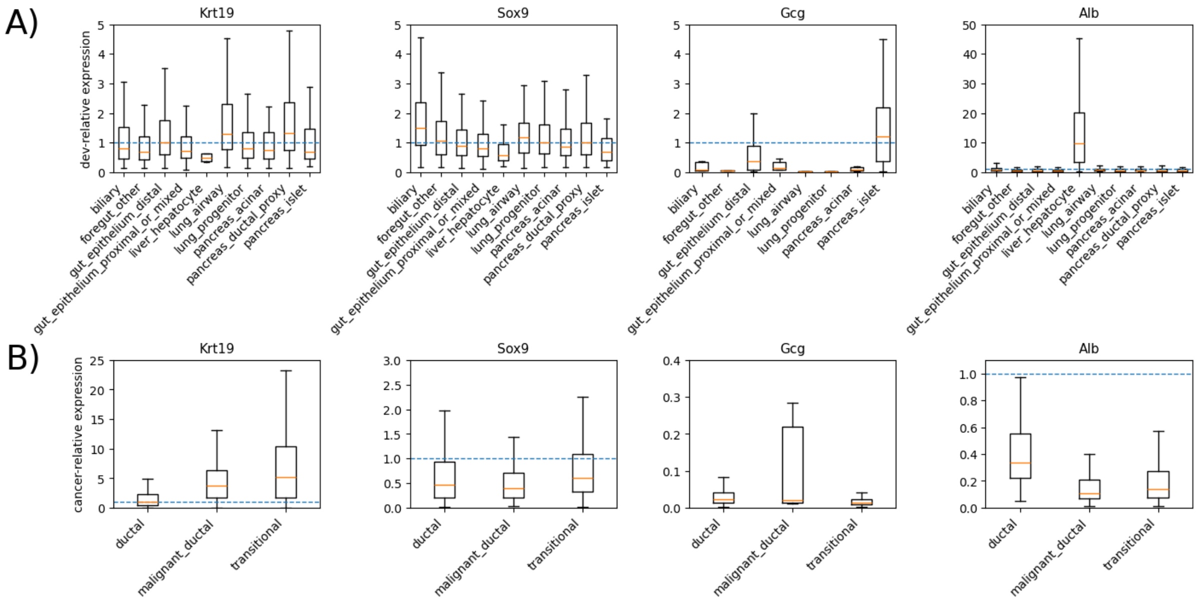


Figure S4: **Validation of dev-relative normalization and cancer-relative deviations across lineage markers** A) Dev-relative expression (dev_rel) of representative lineage markers across developmental endoderm categories. Pan-epithelial markers (Krt19, Sox9) show comparable levels across epithelial tissues, while lineage-restricted genes (Gcg, Alb) remain confined to their appropriate compartments, indicating preservation of developmental structure after normalization. B) Cancer-relative expression (cancer_rel) of the same markers across PDAC cell states. Epithelial markers (Krt19, Sox9) are elevated relative to developmental reference levels, whereas non-pancreatic lineage markers (Gcg, Alb) remain suppressed, demonstrating structured deviations from developmental grammar in PDAC. Dashed lines indicate the developmental baseline defined as a relative expression value of one.

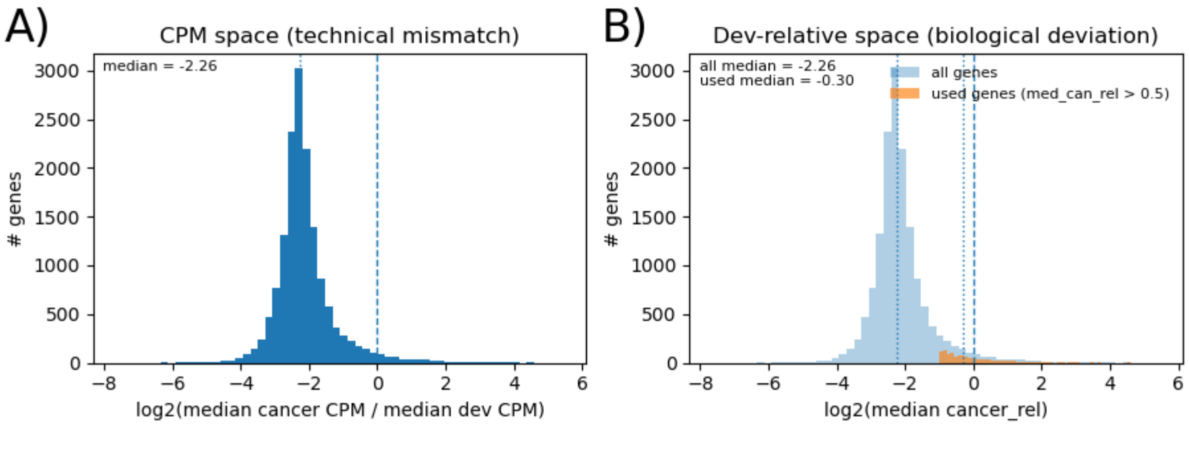


Figure S5: **Validation of dev-relative normalization and cancer-relative deviations across lineage markers** A)*Per*-gene nonzero median expression values compared directly in CPM space show a strong global shift between developmental and PDAC datasets (median $\log_2 \approx$ -2.26), indicating that raw CPM values are not numerically comparable across platforms. B) After dev-relative normalization, *per*-gene median cancer expression reflects structured biological deviation: PDAC underuses most developmental programs but shows near-centered expression for genes active in cancer (median cancer_rel > 0.5; median $\log_2 \approx$ -0.30), consistent with scale alignment within the shared epithelial grammar.

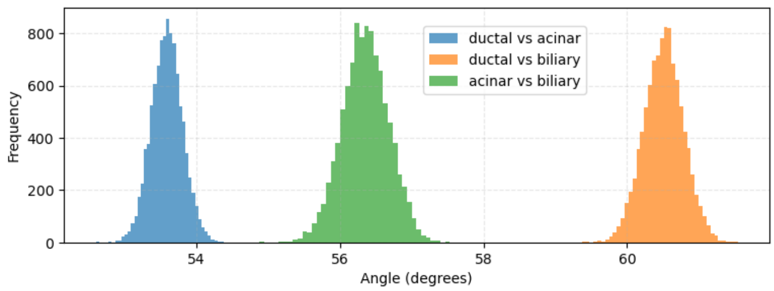


Figure S6: **Stability of developmental lineage directions.** Bootstrap distributions of pairwise angles between centroid-derived developmental lineage vectors from foregut epithelium toward ductal, acinar, and biliary terminal states ($n = 10000$). The three comparisons show narrow distributions centered at approximately 53.6°, 56.3°, and 60.5°, indicating that the lineage-referenced directions are reproducible and geometrically distinct.

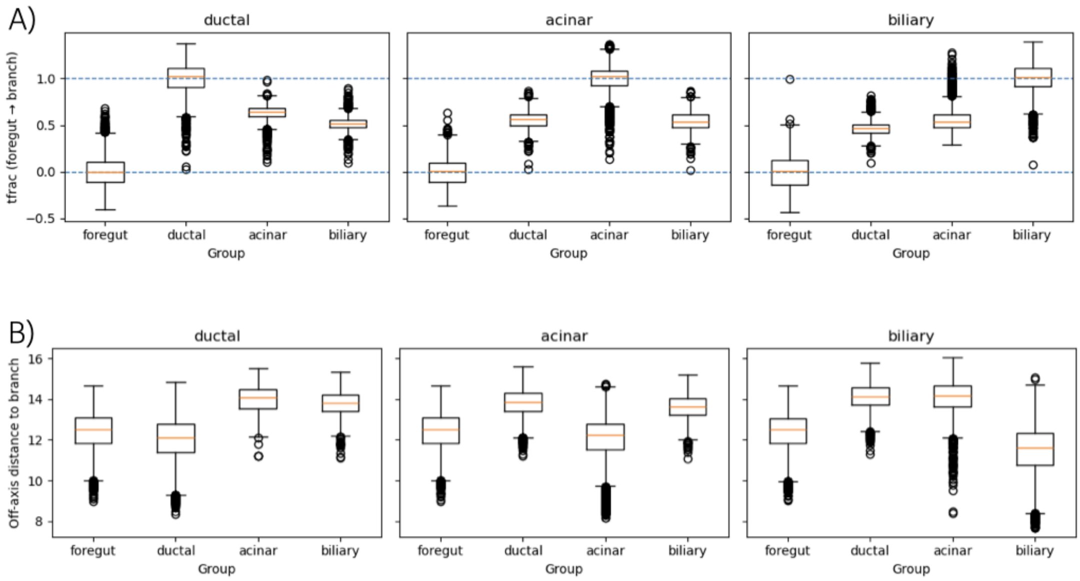


Figure S7: **Developmental cells occupy broad lineage-referenced corridors rather than narrow one-dimensional branch axes.** A) Normalized longitudinal position, $t_{\mathrm{frac}}$, of foregut, ductal, acinar, and biliary cells projected onto each centroid-defined lineage direction. Dashed lines indicate the foregut anchor, $t_{\mathrm{frac}} = 0$, and the corresponding terminal-state position, $t_{\mathrm{frac}} = 1$. For each lineage direction, the matching terminal population is centered near $t_{\mathrm{frac}} = 1$, while non-matching developmental populations also occupy intermediate positions. B) Orthogonal distance, $d_{\perp}$, from each cell to the corresponding lineage axis. Developmental cells are broadly distributed around, rather than tightly confined to, the centroid-to-centroid branch lines.

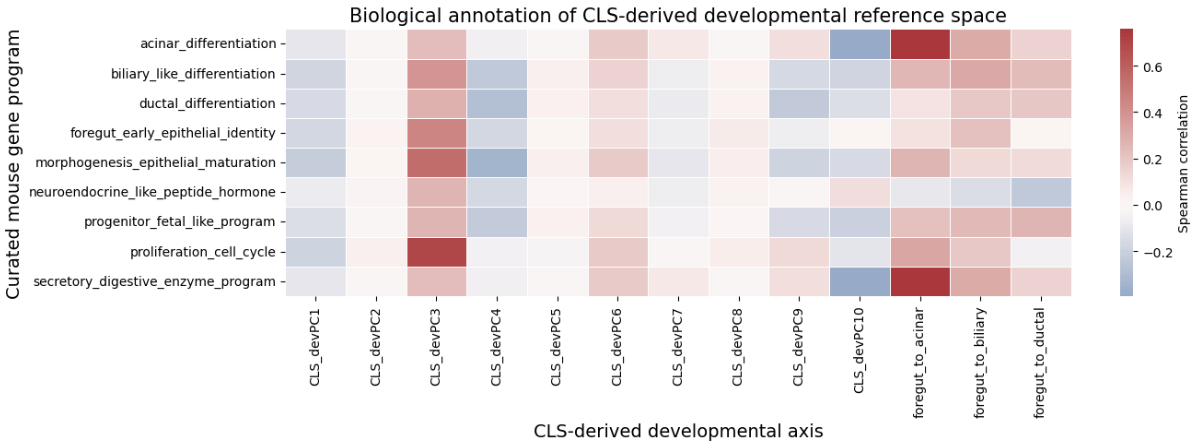


Figure S8: **Biological annotation of CLS-derived developmental reference axes.** Spearman correlations between curated mouse developmental gene program scores and CLS-derived developmental axes. Columns include the first ten principal components (PCs) and centroid-defined lineage directions ($t_i$ values in Eqn. 1). Rows correspond to curated gene programs (`Supplementary note 1`). Positive values indicate that a program increases along the positive direction of an axis, whereas negative values indicate enrichment at the opposite loading. For generic PCs, axis signs are arbitrary and should be interpreted as opposing loadings rather than absolute developmental directions; for centroid-defined lineage projections, positive values indicate movement away from the foregut centroid toward the indicated endpoint. The foregut-to-acinar direction showed the strongest positive association with acinar differentiation and secretory digestive enzyme programs, while `CLS_devPC3` (developmental PC3) was strongly associated with proliferation/cell-cycle and epithelial maturation programs.

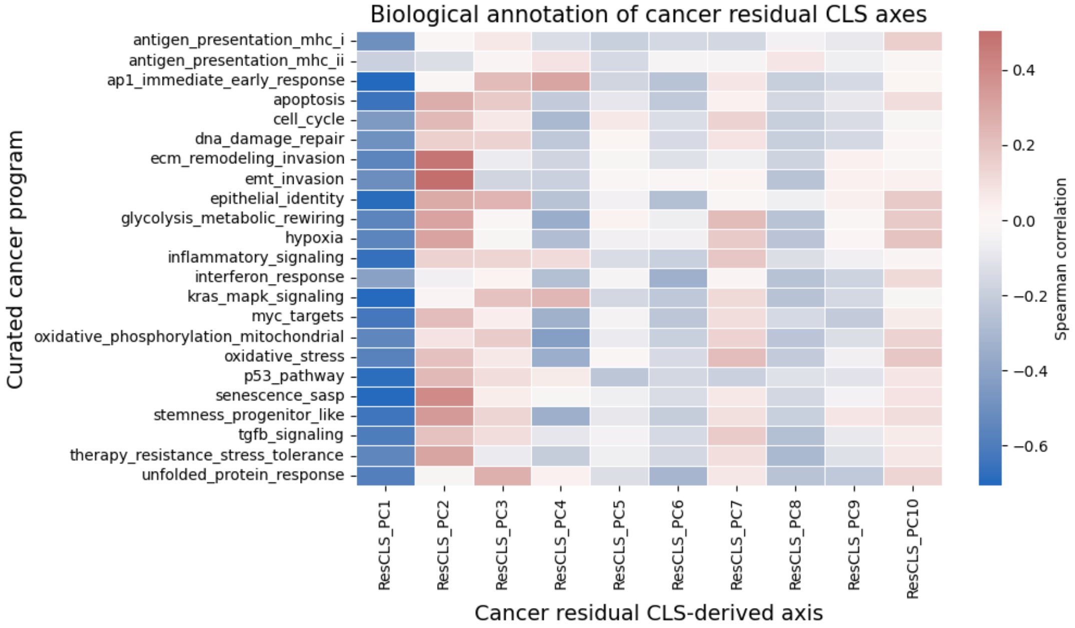


Figure S9: **Biological annotation of residual cancer CLS axes.** Spearman correlations between curated cancer-associated gene program scores and residual cancer CLS axes. Residual axes were computed from the component of malignant CLS variation remaining after reconstruction from the developmental reference basis. Rows correspond to curated cancer programs (`Supplementary note 2`). Columns correspond to the first ten residual cancer PCs. Positive correlations indicate enrichment along the positive loading of each residual axis, whereas negative correlations indicate enrichment at the opposite loading. The residual cancer space contained both residual pancreatic differentiation-like structure and cancer-adaptive programs, including a prominent EMT/ECM invasion axis and an AP-1/immediate early stress-response axis.

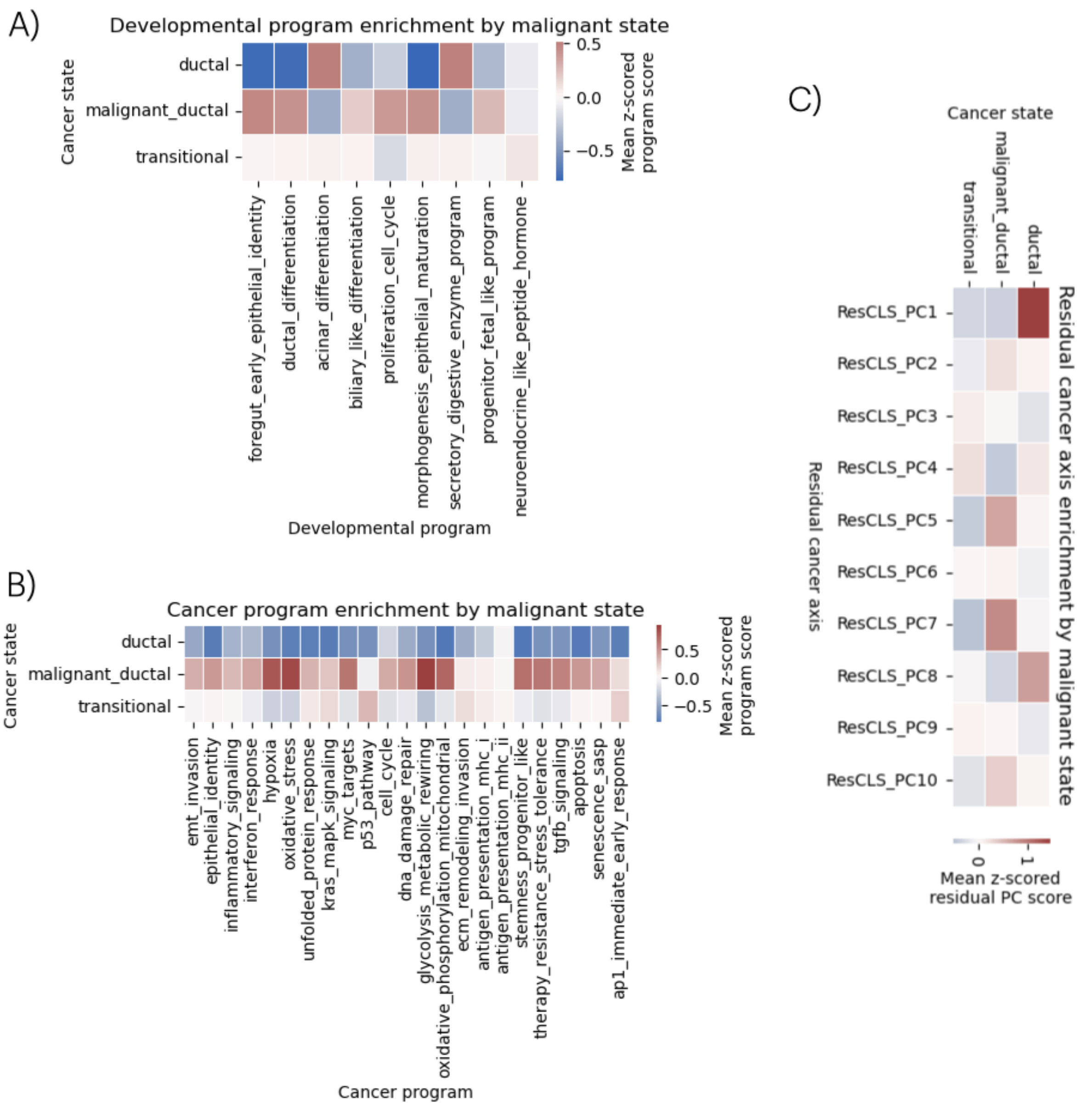


Figure S10: **Malignant state annotation by developmental programs, cancer programs, and residual cancer axes.** Heatmaps show state-level enrichment of residual cancer-associated axes, developmental programs, and cancer-associated programs across malignant states. A) Developmental program enrichment was computed by scoring each developmental program in individual cancer cells, $z$-scoring scores across cancer cells, and averaging within each malignant state. B) Cancer program enrichment was computed analogously using curated cancer-associated programs. C) Residual cancer-axis enrichment was computed as the mean $z$-scored residual PC score for each malignant state. The state initially labeled *ductal* was enriched for acinar differentiation and secretory digestive enzyme programs and showed high positive enrichment along the acinar/secretory residual cancer-associated axis. The *malignant ductal* state showed elevated glycolytic rewiring, oxidative stress, hypoxia, oxidative phosphorylation, stemness/progenitor-like identity, MYC targets, and residual glycolytic/oxidative-stress axes. The *transitional* state showed modest neuroendocrine-like peptide hormone enrichment and modest activation of p53, AP-1/KRAS, ECM remodeling, unfolded protein response, and antigen-presentation programs, supporting its interpretation as a malignant adaptive/transitional state rather than a normal developmental intermediate. Developmental lineage-axis alignment is summarized in Supplementary Table S1 and visualized in Fig.fig3A, main text.

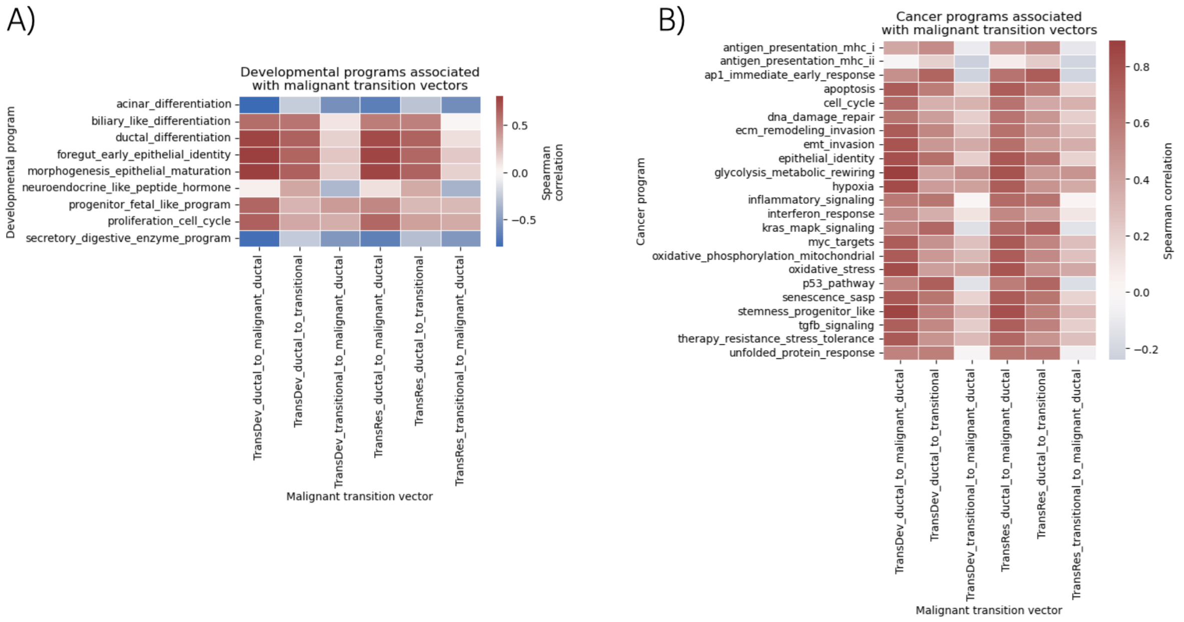


Figure S11: **Biological programs associated with malignant transition vectors.** Heatmaps show Spearman correlations between malignant transition coordinates and curated A) developmental or B) cancer-associated gene program scores. Transition vectors were defined between malignant state centroids in either developmental PC space (‘TransDev’) or residual cancer PC space (TransRes’). Each cancer cell was projected onto each transition vector, and the resulting transition coordinate was correlated with program scores across cancer cells. Positive correlations indicate programs increasing toward the endpoint state, whereas negative correlations indicate programs enriched near the start state. Malignant transition vectors showed loss of acinar differentiation and secretory digestive enzyme programs and gain of ductal/foregut epithelial, progenitor-like, glycolytic, hypoxic, oxidative-stress, MYC, EMT/invasion, AP-1/KRAS, p53, inflammatory, senescence/SASP, unfolded protein response, and proliferation programs. These results indicate that malignant transitions follow cancer-adaptive reprogramming trajectories rather than normal developmental lineage directions.
